# Caveolin assemblies displace one bilayer leaflet to organize and bend membranes

**DOI:** 10.1101/2024.08.28.610209

**Authors:** Milka Doktorova, Sebastian Daum, Tyler R. Reagle, Hannah I. Cannon, Jan Ebenhan, Sarah Neudorf, Bing Han, Satyan Sharma, Peter Kasson, Kandice R. Levental, Kirsten Bacia, Anne K. Kenworthy, Ilya Levental

## Abstract

Caveolin is a monotopic integral membrane protein, widely expressed in metazoa and responsible for constructing enigmatic membrane invaginations known as caveolae. Recently, the high-resolution structure of a purified human caveolin assembly, the CAV1-8S complex, revealed a unique organization of 11 protomers arranged in a tightly packed, radially symmetric spiral disc. One face and the outer rim of this disc are hydrophobic, suggesting that the complex incorporates into membranes by displacing hundreds of lipids from one leaflet. The feasibility of this unique molecular architecture and its biophysical and functional consequences are currently unknown. Using Langmuir film balance measurements, we find that CAV1-8S is highly surface active, intercalating into lipid monolayers of various compositions. CAV1-8S can also incorporate into preformed bilayers, but only upon removal of phospholipids from the outer-facing leaflet. Atomistic and coarse-grained simulations of biomimetic bilayers support this ‘leaflet replacement’ model and also reveal that CAV1-8S accumulates 40−70 cholesterol molecules into a disordered monolayer between the complex and its distal lipid leaflet. We find that CAV1-8S preferentially associates with positively curved membrane surfaces due to its influence on the conformations of distal leaflet lipids, and that these effects laterally sort lipids. Large-scale simulations of multiple caveolin assemblies confirmed their association with large, positively curved membrane morphologies consistent with the shape of caveolae. Further, association with curved membranes regulates the exposure of caveolin residues implicated in protein-protein interactions. Altogether, the unique structure of CAV1-8S imparts unusual modes of membrane interaction with implications for membrane organization, morphology, and physiology.

**STATEMENT OF SIGNIFICANCE:** Caveolae are membrane invaginations heavily implicated in cellular physiology and disease; however, how their unique shape and function are produced remains enigmatic. Here, following on recent characterization of the unusual structure of the CAV1-8S oligomer, we examine the molecular details of its interactions with its surrounding lipid membrane using simulations and reconstitution experiments. We describe a novel mode of membrane interaction−which we term ‘leaflet replacement’−for the CAV1-8S complex that has not previously been observed for any other protein. The biophysical consequences of this unique molecular organization provide mechanistic insights into the functions and organization of caveolae in cells.

## INTRODUCTION

Membrane organization underlies many cellular functions, including signal transduction, subcellular trafficking, and host-pathogen interactions (1). This organization includes lateral lipid domains (e.g. lipid rafts) (2), membrane protein clusters (3, 4), membrane curvatures (5-7), and combinations of these (8-10). A prominent cellular feature that incorporates several of these aspects are caveolae, 50-100 nm flask-shaped plasma membrane invaginations that sense, organize, and relay signals in a wide range of cell types (11-15). Caveolae require expression of unusual monotopic integral membrane proteins called caveolins (in humans, three family members CAV1-3), which assemble into high-order oligomers that serve as the fundamental caveolae building blocks (16, 17). A family of peripheral membrane proteins, the cavins, contributes to caveolar biogenesis together with specific lipids and accessory proteins such as pacsin2 and EHD2 (18-21, 22)). While a detailed understanding of how cavins associate with lipids has emerged (23-26), much less is known about how caveolins interact with membranes and contribute to the formation of caveolae (11). For example, whereas caveolin-rich membrane fractions are enriched in cholesterol and sphingolipids and the structural integrity of caveolae is highly sensitive to cellular cholesterol levels (27), the molecular mechanisms underlying lipid organization by caveolins are not understood (21). Similarly, the often-hypothesized compositional, structural and functional connections between caveolae and lipid rafts (28) remain poorly resolved.

Caveolin has been associated with various extents of membrane curvature: deep invaginations (which can be fully internalized (19, 29)), shallow ‘dolines’ (30), or even fully flattened membrane domains (31, 32). However, how caveolin-lipid interactions contribute to its preference for curved regions and whether caveolin itself is a driver of curvature is not known. One pertinent question is how the unusual structure and oligomeric assembly of caveolins is connected to the shape and functions of caveolae. A long-standing hypothesis suggests that caveolins induce membrane curvature by inserting a hydrophobic hairpin into the cytoplasmic leaflet (20, 33-36). An alternative model prompted by recent structural evidence suggests that caveolae are shaped by a flexible network of cavin-1 that surrounds membrane-embedded oligomeric discs of caveolin (37). The molecular architecture of such oligomeric discs, referred to as 8S complexes, was recently determined for human caveolin-1 (CAV1-8S) (38). The cryoEM structure of CAV1-8S consists of 11 copies of CAV1 assembled into spirals that form a tightly packed disc with a diameter of ∼14 nm (Fig 1A, S1A−C). One face of the disc is flat, while the other contains a raised rim and a protruding central beta-barrel. Strikingly, the flat disc face is almost exclusively hydrophobic (Fig. 1A), suggesting intimate association with the hydrophobic core of the membrane (38). The outer rim of the disc (∼3 nm deep) is also largely hydrophobic. This unique architecture suggests a model in which the CAV1-8S complex is fully embedded in the cytoplasmic leaflet of the plasma membrane, potentially displacing a large patch of lipids, with its membrane-facing surface fully solvated by the lipid chains of the distal, extracellular monolayer (21, 38) (Fig. 1A). Whether this arrangement is feasible and how it affects lipid organization and membrane curvature are currently unknown.

**Figure 1.**
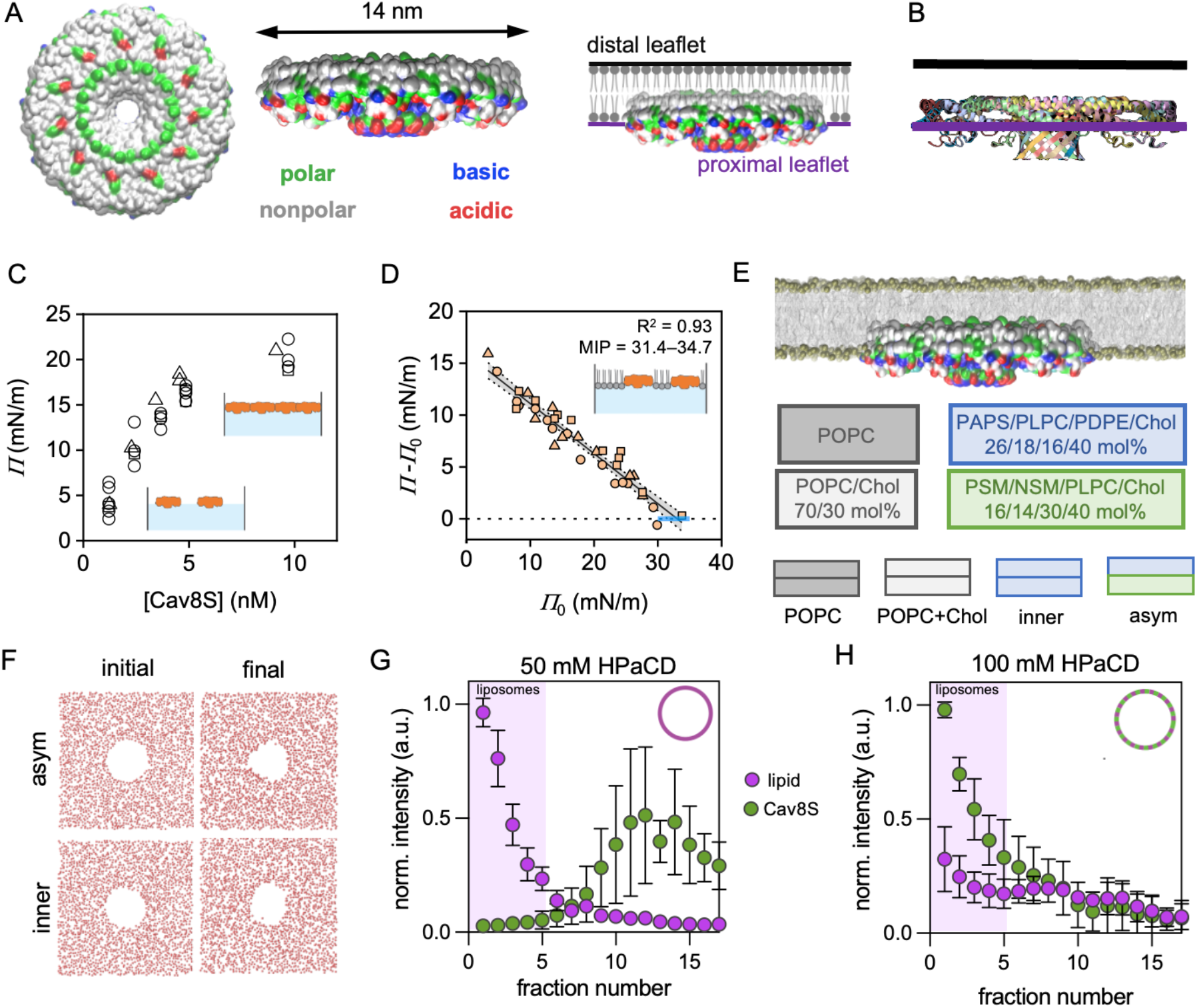
CAV1-8S incorporates into a single leaflet by displacing many lipids. (A) Surface representation of the membrane-facing surface (left) and side rim (middle) of CAV1-8S with residues colored according to hydrophobicity and charge, as indicated. The hydrophobic face and rim suggest that the complex can embed into the bilayer by displacing a large patch of lipids from the protein-proximal leaflet (right). (B) Positioning of CAV1-8S in a generic bilayer predicted by the PPM server. Colors represent individual protomers of the complex. (C) Changes in surface pressure caused by addition of CAV1-8S to the subphase under a lipid monolayer. (D) CAV1-8S inserts into highly compressed POPC monolayers up to 30−35 mN/m. Plotted is the difference in pressure upon adding CAV1-8S to POPC at various surface pressures at the water-air interface (see Fig. S2B). The x-intercept indicates the maximum monolayer insertion pressure (MIP), which for CAV1-8S coincides with the equivalence pressure (EQP) at which the packing of lipid monolayers is approximately equivalent to that of lipid bilayers. Symbols in C-D represent three independent protein preparations. (E) Illustration of the initial CAV1-8S placement in the simulated membranes and their leaflet lipid compositions. (F) Phospholipid distribution in the CAV1-8S-proximal leaflet (viewed from above the leaflet) at the beginning (left) and end (right) of the asym (top) and inner (bottom) lipid bilayers. Red dots represent positions of lipid phosphate groups in the protein-proximal leaflet. (G) CAV1-8S does not incorporate into large unilamellar vesicles (LUVs) in the presence of 50 mM HPaCD, but (H) does incorporate at 100 mM HPaCD, which extracts sufficient lipids to induce the onset of membrane disruption (Fig. S3E). Mean ± st. dev. from 3 independent experiments.

Here, combining experiments and simulations, we examined how CAV1-8S interacts with and organizes lipid membranes. We validated the hypothesis that CAV1-8S can fully embed into bilayers by displacing phospholipids from its proximal leaflet and investigated the consequences of this peculiar arrangement on lipid sorting, cholesterol dynamics, membrane curvature, and accessibility of protein-interaction residues. We find that the distinctive mode of membrane insertion of CAV1-8S recruits cholesterol, prefers positively curved surfaces, and laterally organizes lipids. These insights begin to define the structural mechanisms underlying the physiology of caveolae in cells.

## RESULTS

### CAV1-8S displaces phospholipids to fully embed in its proximal leaflet

Previous investigations into caveolin’s membrane insertion and curvature have focused on monomers or low-order assemblies (30, 39-45), whereas the functional units of caveolae are believed to be higher-order complexes, represented by CAV1-8S. The dynamics of CAV1-8S in solution and on implicit membrane vesicles were recently investigated using molecular dynamics (46); however, how the complex is embedded into and remodels lipid membranes remains unknown. The recently resolved CAV1-8S structure suggests a unique mode of interaction with the lipid matrix: the large, tightly packed hydrophobic surface and its hydrophobic rim (Fig. 1A) may embed deeply in the bilayer by displacing a large patch of lipids from the protein-proximal leaflet (17, 21, 38, 47). To evaluate this possibility, we first used the Positioning of Proteins in Membranes (PPM) web server (48, 49) to model the protein’s position within a generic membrane. PPM predicted that CAV1-8S embeds fully into one of the membrane leaflets (Fig. 1B).

To experimentally evaluate the surface activity and lipid monolayer incorporation of CAV1-8S, we performed film balance experiments on a Langmuir monolayer, which detects changes in surface pressure resulting from the partitioning of amphiphilic molecules to the aqueous-air interface. Injection of purified CAV1-8S (9.7 nM) into the aqueous subphase produced a rapid increase in surface pressure that equilibrated at an elevated pressure, indicating high surface activity (i.e. amphiphilicity) of purified CAV1-8S (Fig. S2A). The equilibrated surface pressure (related to adsorption to the air-water interface) of CAV1-8S was concentration-dependent, approaching saturation at 4−6 nM (Fig. 1C). This behavior indicates that the protein is highly surface active and can generate a dense monolayer at the air-water interface, even in the absence of lipids. We used these measurements to estimate the surface area per CAV1-8S complex at the air-water interface: assuming that at the onset of surface saturation (∼4 nM protein injected into the subphase) almost all protein partitions to the interface, the calculated area for a single complex is ∼113 nm^2^, which is in good correspondence with the area of the flat, hydrophobic face of the complex estimated from cryoEM (∼154 nm^2^). While there are significant caveats to these assumptions (some protein likely lost to the Teflon surfaces, incomplete partitioning to the interface), these observations suggest that CAV1-8S strongly prefers the aqueous-hydrophobic interface, where it lays flat with its hydrophobic face exposed to air (Fig 1C, inset).

Given the high surface activity of CAV1-8S, we tested its propensity to insert into lipid monolayers by determining its maximum insertion pressure (MIP) (50) (Fig. 1D). For these measurements, monolayers of lipids at various surface pressures (*Π*_0_) are produced by spreading various lipid amounts from organic solution onto the air-water interface. After solvent evaporation, CAV1-8S is injected into the aqueous subphase under the monolayer, which leads to an increase in pressure from *Π*_0_ to *Π* (i.e. *ΔΠ*) as the protein inserts into the lipid monolayer (Fig. S2B). The magnitude of *ΔΠ* represents the protein’s ability to partition into the compressed lipid monolayer; linear regression of *ΔΠ* as a function of *Π*_0_ yields the protein’s MIP (see Fig. 1D). We found that CAV1-8S partitioned strongly into lipid monolayers of various compositions (POPC only, POPC + 25 mol% Chol, equimolar mixture of POPC, POPS and POPE), even at high lipid density (Fig. 1D, S2C−D). The MIP values for CAV1-8S were 33.0, 37.8 and 41.1 mN/m in POPC, POPC/Chol and POPC/POPS/POPE, respectively (Table S1). These values are all within or above the range of the bilayer Equivalence Pressure (EQP), where the monolayer lipid packing density equals that of a self-assembled lipid bilayer (30−35 mN/m (50)), indicating that the CAV1-8S complex can embed into tightly packed lipid environments. In contrast, a control protein that is not expected to partition efficiently into membranes (Sar1-ΔN, the small GTPase Sar1 lacking its N-terminal amphipathic helix (51)), had a MIP <25 mN/m, i.e. below the EQP and therefore consistent with inefficient monolayer insertion (Fig. S2E, Table S1). These experiments confirm that CAV1-8S can embed into lipid monolayers even at the lipid packing densities of biological membranes.

To obtain molecular insight into the interaction of CAV1-8S with lipid bilayers, we performed molecular dynamics simulations. Because the possible membrane remodeling and lipid sorting by CAV1-8S requires simulation systems that are too large to effectively simulate with all-atom models, our initial experiments relied on the MARTINI coarse-grained model, which successfully recapitulates key properties of protein-membrane systems on extended time and length scales (52). We constructed large bilayers (45×45×27 nm, ∼7000 lipids) of increasing complexity and biological relevance: pure ***POPC*** as a minimal model of a biomembrane, ***POPC+Chol*** to include the highly abundant and structurally important cholesterol, a 4-component mixture representing the inner leaflet of the mammalian PM (***inner***), and an asymmetric model representing the complex, asymmetric lipid distribution of the PM (***asym***) (Fig 1E, compositional details in Table S2). To test the ‘leaflet replacement’ model we inserted CAV1-8S according to the positioning suggested by the PPM prediction (Fig. 1B), which required removal of ∼250 lipids (<4% of the bilayer) from the protein-proximal monolayer to maintain the lipid packing density of the pure lipid bilayer. Across these protein-lipid systems simulated for tens of microseconds, CAV1-8S maintained its deeply embedded orientation, effectively displacing all phospholipids from underneath the complex in its proximal leaflet, while distal leaflet lipids solvated its flat hydrophobic surface (Fig. 1F, S1D−E). These results demonstrate that the ‘leaflet replacement’ model of CAV1-8S membrane intercalation represents a physically feasible and stable arrangement.

To experimentally verify the computational prediction that removing lipids from a membrane leaflet allows stable CAV1-8S incorporation, we investigated the incorporation of purified CAV1-8S-mVenus (labeled at the C-terminus, which preserves the oligomeric structure (53)) into preformed lipid vesicles (extruded ∼100 nm liposomes). We measured membrane association by co-flotation, wherein liposomes and membrane-incorporated proteins float to the top of sucrose gradients after centrifugation while proteins not associated with membranes remain at the bottom of the gradient. CAV1-8S did not incorporate into undisturbed liposomes, with minimal fluorescence from the protein (i.e. mVenus) found in the top, liposome-containing fractions (Fig. S3A). This was an expected result, since insertion of such a large complex into a tightly packed bilayer would likely have major kinetic barriers and also introduce a major area imbalance between the two leaflets. According to our model (Fig. 1A), both effects could be alleviated by removal of phospholipids from the leaflet hosting the complex. To test whether removal of lipids from the bilayer would facilitate CAV1-8S incorporation, we used hydroxypropyl-α-cyclodextrin (HPaCD) to extract outer leaflet phospholipids, as previously described (54, 55). Titration with HPaCD into lipid-only liposomes showed that ∼100 mM HPaCD led to the onset of liposome solubilization, a consequence of excess lipid removal from one leaflet that precludes stable bilayers (54, 56) (Fig S3E). Notably, the inclusion of CAV1-8S prevented this solubilization, suggesting that the protein complex protects liposomes from stresses induced by extraction of lipids from one leaflet (Fig 1H and S3C-D). The mechanism of this protection is likely its incorporation into the outer liposome leaflet, evidenced by its presence in the top, liposome-containing fractions only under HPaCD concentrations that induce bilayer stress (Fig 1G-H, S3A-D). The liposomes formed by CAV1-8S incorporation after lipid extraction maintained similar size profiles as undisturbed controls (Fig S3F). These observations were unique to CAV1-8S, as neither a soluble (StayGold), peripheral (SNAP25), or transmembrane (Syntaxin-1A) protein incorporated efficiently into preformed liposomes either in unperturbed or HPaCD-treated conditions (Fig S4). These results are consistent with the hypothesis that CAV1-8S insertion into a membrane requires displacement of large numbers of lipids, supporting the ‘leaflet replacement’ model.

### CAV1-8S associates with curved membrane regions

The large coarse-grained simulations allowed us to examine changes in membrane morphology associated with CAV1-8S. In both single-component POPC bilayers and cholesterol-containing bilayers, we observed regions of membrane curvature near CAV1-8S, with the distal monolayer curving towards the complex (Fig. 2A, S5) (we refer to this curvature as ‘positive’, according to convention in cells, where positive curvature is towards the cytoplasm). In cholesterol-containing membranes, these curvatures were generally less sharp and spread over a larger area (Fig. 2B, S5). While highly curved membrane regions were consistently colocalized with CAV1-8S across all simulated membranes, it was possible that the construction of the systems may contribute to this effect by introducing asymmetry in leaflet surface areas. Namely, if a large discrepancy exists between the surface area of the lipids removed to insert CAV1-8S into a leaflet and the area of the inserted protein, the bilayer will tend to curve towards the leaflet with the larger area. To control for this possibility, we constructed a large bilayer model with two CAV1-8S complexes embedded in opposing leaflets, with each displacing 250 lipids (Fig. 2C). This bilayer was symmetric, precluding leaflet area differences; however, it also gradually developed a sharp buckling deformation with the two protein complexes both localizing to the two regions of largest positive (i.e. towards the complex) curvatures (Fig. 2C).

**Figure 2.**
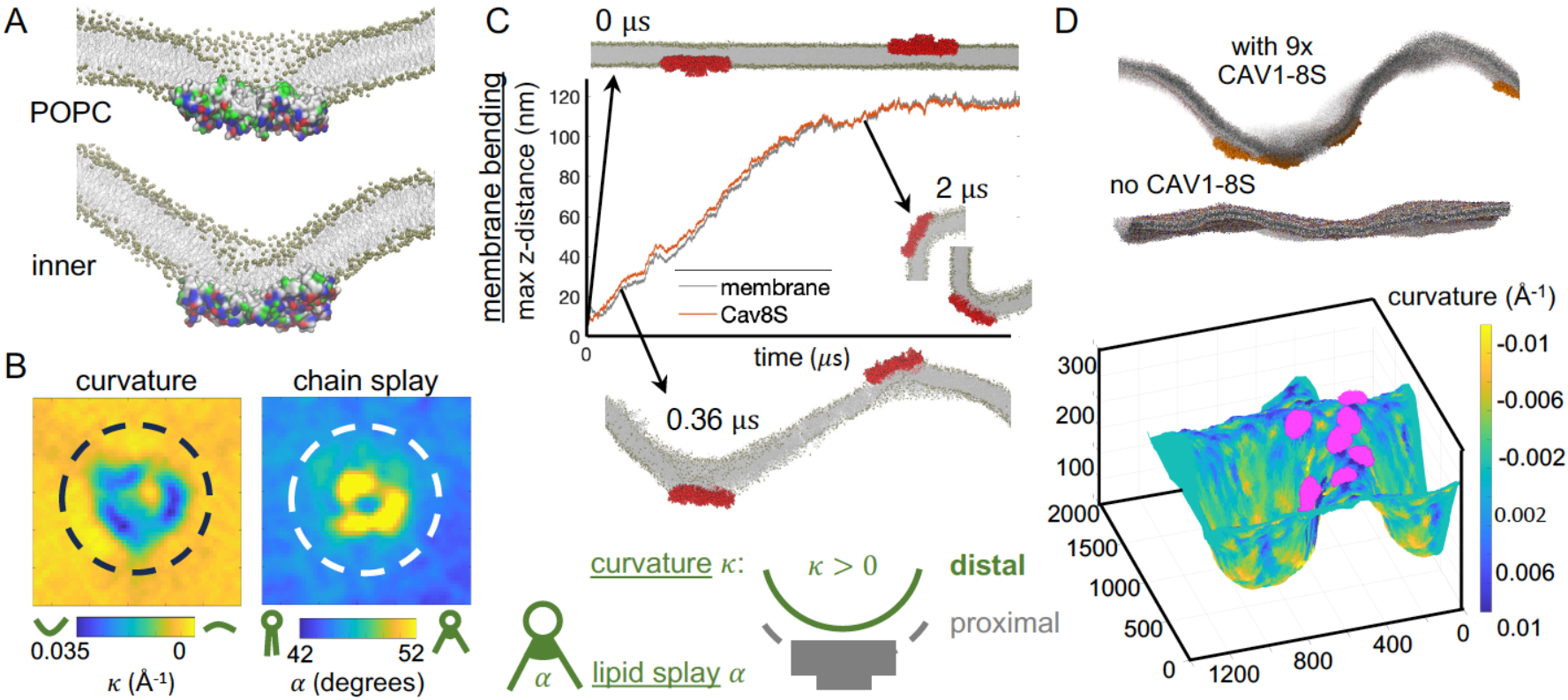
The caveolin complex partitions to membrane regions with high curvature. (A) Snapshots of simulations with POPC (top) and inner mixture (bottom). Lipid phosphates shown as tan spheres, lipid tails as gray lines, and protein is in surface representation with residues colored as in Fig. 1A. CAV1-8S shown in cross-section to show arrangement of distal leaflet lipids. (B) Maps of the local curvature κ around CAV1-8S (165×165 Å centered on the complex) in the inner bilayer mixture (positive is towards the protein complex), and the corresponding lipid chain splay, i.e. the angle α between the two lipid acyl chains. Both properties are calculated from the lipids in the distal leaflet, as schematically illustrated on the right; the dotted circles represent outlines of CAV1-8S. (C) Time evolution of the symmetric inner bilayer mixture with two CAV1-8S complexes embedded in the two opposing bilayer leaflets. Plotted are the vertical distances (i.e. the z-direction, normal to the initial bilayer axis) between maximal and minimal lipid headgroup positions (membrane, gray) or caveolin atom positions (red) as a function of simulation time. Representative snapshots at 0, 0.36 and 2 μs are shown for comparison (CAV1-8S complexes in red). (D) Representative snapshots from a slice of a membrane with or without 9 CAV1-8S (shown in orange). (bottom) Surface rendering with colors representing distal leaflet curvature. CAV1-8S (magenta) is primarily associated with regions of positive curvature. The curvature map is flipped relative to the snapshot for visualization.

To understand the mechanism underlying these observations, we turned to recent theoretical insights that modeled curvature generation by the flat discs of caveolin (57). This model is based on differences in interaction energy of distal leaflet lipid acyl chains with the flat face of CAV1-8S versus with lipids from the proximal leaflet. The underlying assumption is that lipid acyl chains prefer to interact with one another across the bilayer midplane rather than with the protein. In this scenario, distal leaflet lipids in contact with CAV1-8S would splay out their acyl chains to minimize total lipid contacts with the protein complex. Splaying of the chains changes the shape of these lipids towards a more conical conformation, which tends to promote membrane curvature. Indeed, lipid splay was notably increased for distal leaflet lipids above CAV1-8S in both coarse-grained (Fig. 2B & S5) and all-atom simulations (Fig. S6), which also both showed regions of localized positive membrane curvature of the exoplasmic leaflet. Altogether, these results support a mechanistic explanation for the curvature preference of the caveolin complex.

Individual 14 nm discs are likely insufficient to form 100-nm diameter caveolae (57, 58). To begin to understand how multiple caveolin complexes may assemble on membranes, we simulated large coarse-grained (CG) systems with 6 or 9 CAV1-8S assemblies (12-35 μs per replica, Table S2). In these simulations, all complexes again associated with positively curved membrane regions (Fig. 2D). This tendency appeared to be cooperative and via long range effects, as multiple complexes assembled onto large, curved ridges despite minimal direct interactions between the assemblies.

An important consideration for large CG membrane models is their tendency to spontaneously bend. While we observed significant membrane curvature in all systems with CAV1-8S, similar simulation conditions have been reported to produce unphysically large membrane undulations even in the absence of protein (59). We also observed significant undulation of the large CG membranes in control simulations without CAV1-8S, though in most cases their magnitudes were notably smaller than the curvatures associated with CAV1-8S (Fig. 2D). Ultimately, while our results do not conclusively attribute the changes in bilayer shape to CAV1-8S, they clearly demonstrate that CAV1-8S associates with positively curved membrane regions, consistent with its ability to induce invaginating membrane domains in cellular plasma membranes. Further, the effect of the complex on lipid conformation provides a mechanistic explanation for curvature generation.

### Cholesterol accumulates underneath CAV1-8S

To interrogate lipid rearrangements resulting from the unusual positioning of CAV1-8S within the membrane, we analyzed the lipid compositions and conformations under the complex in our CG simulations. For readability and related to the native localization of caveolin in the cytoplasmic leaflet, we define the lipid leaflet containing CAV1-8S as ‘cytoplasmic’ and the opposite leaflet as ‘exoplasmic’. While CAV1-8S stably displaced phospholipids from the ‘cytoplasmic’ leaflet across lipid compositions (Fig 1F, S1D−E), cholesterol (if present) rapidly and specifically accumulated directly underneath the protein (Fig. 3A−B). This accumulation produced an unusual configuration: an ‘exoplasmic’ leaflet composed of phospholipids (PLs) and cholesterol, and a cytoplasmic leaflet containing CAV1-8S with tens of cholesterol molecules solvating its large hydrophobic surface and separating the complex from the exoplasmic leaflet (Fig. 3A). The abundance of cholesterol recruited by CAV1-8S varied with lipid composition: in POPC+30 mol% Chol, ∼40 cholesterol molecules were accumulated, while in the more complex mixtures with 40 mol% Chol, up to ∼70 were recruited (Fig. 3B). Notably, the orientations of these cholesterol molecules were quite distinct from cholesterol in the exoplasmic leaflet (or the bulk cytoplasmic leaflet): they were more disordered, as shown by the broader distribution of cholesterol tilt angles (Fig. S7). We also observed specific accumulation of cholesterol molecules within the hydrophobic core of the CAV1-8S beta-barrel, which is contiguous with the flat, hydrophobic membrane-facing surface (Fig. 3C). Up to 6 Chol molecules accumulated inside the barrel, with their headgroups solvated by the water on the cytoplasmic side of the complex (Fig. 3C). We confirmed these observations in all-atom simulations of the complex embedded in a POPC/Chol bilayer (Fig S8). Similar to the coarse-grained trajectories, Chol molecules accumulated under the CAV1-8S complex, where they adopted a wide distribution of tilt angles (Fig. S8B). Some of these cholesterols then flipped and traversed through the β-barrel, with their headgroups pointing out of the barrel into the cytosol. While the limited timescale of these unbiased trajectories (1 *μ*s) prevents statistically robust quantitation of Chol accumulation, the observations are fully consistent with recent metadynamics simulations (58) and our CG results, and confirm the favorable interaction of Chol with the hydrophobic interior of the β-barrel. Thus, each CAV1-8S not only displaces PLs from the cytoplasmic leaflet but also recruits and sequesters tens of cholesterol molecules into a disordered, pure-cholesterol monolayer, as well as several cholesterol molecules within its β-barrel.

**Figure 3.**
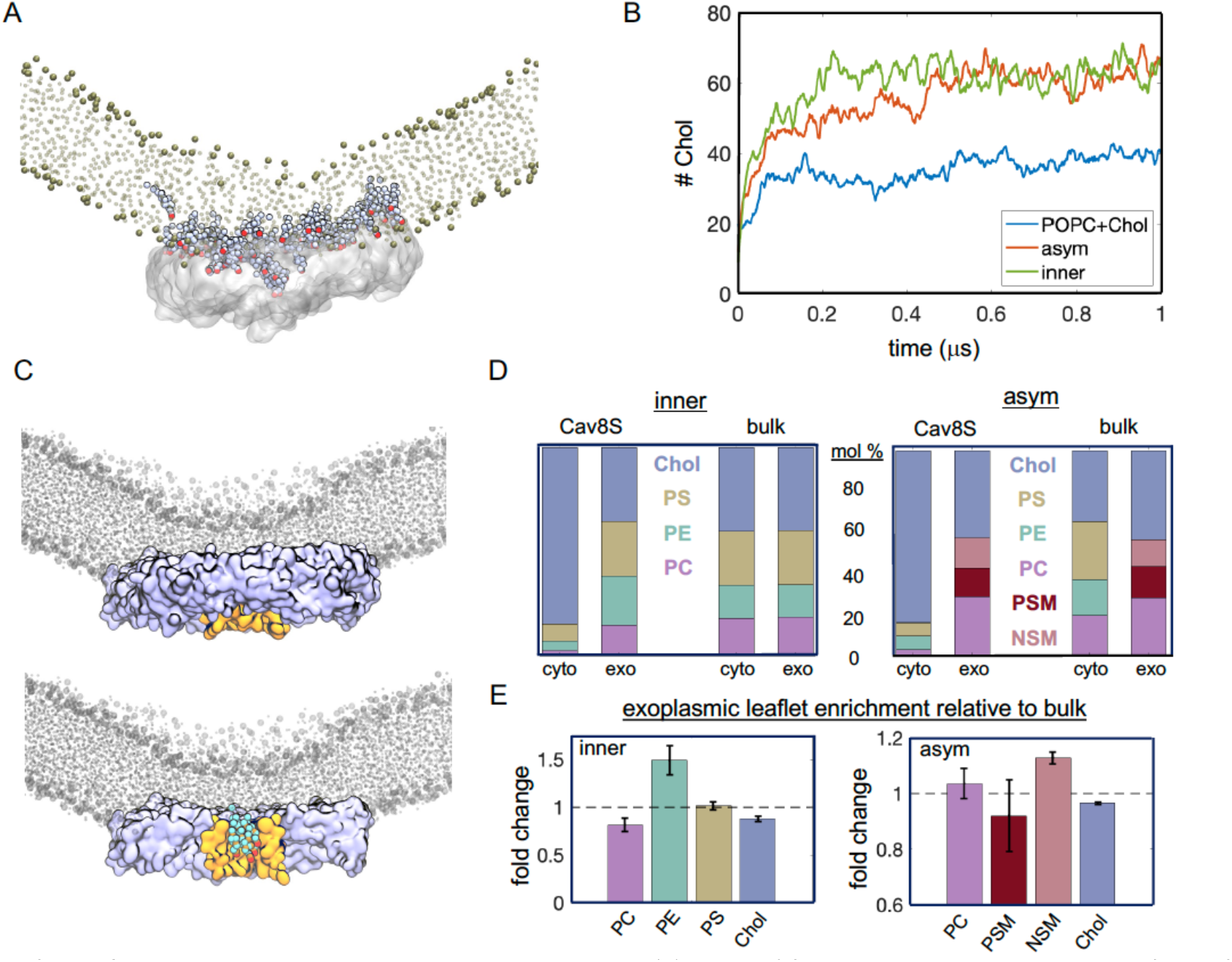
CAV1-8S recruits cholesterol and organizes phospholipids. (A) A layer of Chol molecules is recruited between CAV1-8S and the distal leaflet. Shown is a snapshot of an inner bilayer simulation. Protein is shown in white surface representation, lipid headgroups and chains shown as tan spheres and in Licorice representation, respectively. Chol molecules in contact with the complex are shown in VdW representation with -OH headgroups in red. (B) Time course of Chol accumulation (number of molecules under CAV1-8S) in the cytoplasmic leaflet during the first 1 *μ*s of simulation. (C) Rendering from the end of a POPC+Chol simulation in which the central beta barrel of CAV1-8S was maintained (via constraints), showing 5 Chol molecules recruited within the barrel (full side view of the complex on top, cross-section on bottom). POPC lipids in gray, protein is a blue surface with the beta barrel in yellow. The 5 Chol molecules are in cyan with headgroups in red. (D) Lipid and Chol compositions of the cytoplasmic and exoplasmic leaflets under CAV1-8S and in the bulk membrane in the inner bilayer mixture (left) and the asymmetric PM model (right). A lipid was considered under the complex if its phosphate bead was within 2 Å in the *x* and *y* dimensions of any CAV1 bead (Methods). Thus, the cytoplasmic leaflet phospholipids at the periphery of the complex were also counted, though these are a minor fraction. (E) Enrichment (>1) or depletion (<1) of lipids in the exoplasmic leaflet under CAV1-8S, shown as fold change relative to the mol% in bulk of same leaflet.

### CAV1-8S induces lateral lipid reorganization

Despite the prominent enrichment of Chol in the cytoplasmic leaflet under CAV1-8S, Chol concentration in the complex-facing exoplasmic leaflet was similar to the bulk exoplasmic leaflet (Fig 3D). In contrast, we observed significant phospholipid sorting in that leaflet: e.g., in the ***inner*** model (i.e. symmetric bilayer representative of PM inner leaflet lipids), PE was enriched in the ‘exoplasmic’ leaflet facing CAV1-8S while PC was depleted (Fig. 3D−E). These trends likely reflect the curvatures associated with CAV1-8S (Fig. 2), with cone-shaped PE being preferentially recruited to the region of high curvature, consistent with the lipid-splay explanation above (Fig 2B). While this specific effect is likely not biologically relevant−PE is not expected to be in the exoplasmic PM leaflet−it demonstrates the capacity for lipid sorting due to CAV1-8S-associated curvature. In the ***asym*** model representing the complex asymmetric PM bilayer, we observed accumulation of long-chain sphingomyelin (NSM) in the exoplasmic leaflet facing CAV1-8S. This effect was surprisingly selective for SM with longer acyl chains, as the shorter species (palmitoyl SM) was slightly depleted from the region in contact with CAV1-8S (Fig 3E). We speculate that this effect is related to the ability of longer acyl chains to reach across the bilayer to solvate the protein surface and/or interact with the Chol layer in the cytoplasmic leaflet. While detailed analysis of such interactions would require higher resolution models, the general observations are consistent with extensive experimental reports of enrichment of raft-associated lipids like SM and Chol in caveolae (21, 29, 60, 61).

### Palmitoylation attracts and orders cholesterol under CAV1-8S

Each CAV1 monomer contains 3 cysteine residues that are substrates for S-palmitoylation (62). Thus, CAV1-8S can potentially bear as many as 33 palmitate modifications. To determine the effects of palmitoylation on CAV1-8S-membrane interactions, we constructed a model of the complex with all 33 cysteine residues replaced by their palmitoylated analogs. This palmitoylated CAV1-8S recruited a Chol layer even more efficiently than non-palmitoylated, recruiting ∼60% more Chol molecules (Fig. 4A). These Chol molecules were also markedly more ordered relative to those recruited by the non-palmitoylated complex, revealed by the tighter distribution of tilt angles and an average more similar to that of bulk cholesterol (gray bars in Fig. 4B). The protein-coupled palmitoyl chains themselves remained largely disordered, adopting random orientations not aligned with the bilayer normal, quite unlike lipid-coupled palmitates (Fig. S9). Lower order of protein-attached lipid anchors has been previously reported (63, 64) and for CAV1-8S, likely arises from incorporation of the anchors into the relatively thin and disordered cytoplasmic cholesterol layer. Interestingly, the presence of palmitoyl chains also appeared to laterally expand the curvature footprint of CAV1-8S (Fig. 4C). These results indicate that palmitoylation of CAV1-8S enhances Chol recruitment and order, in agreement with recent all-atom simulations of the palmitoylated complex (58).

**Figure 4.**
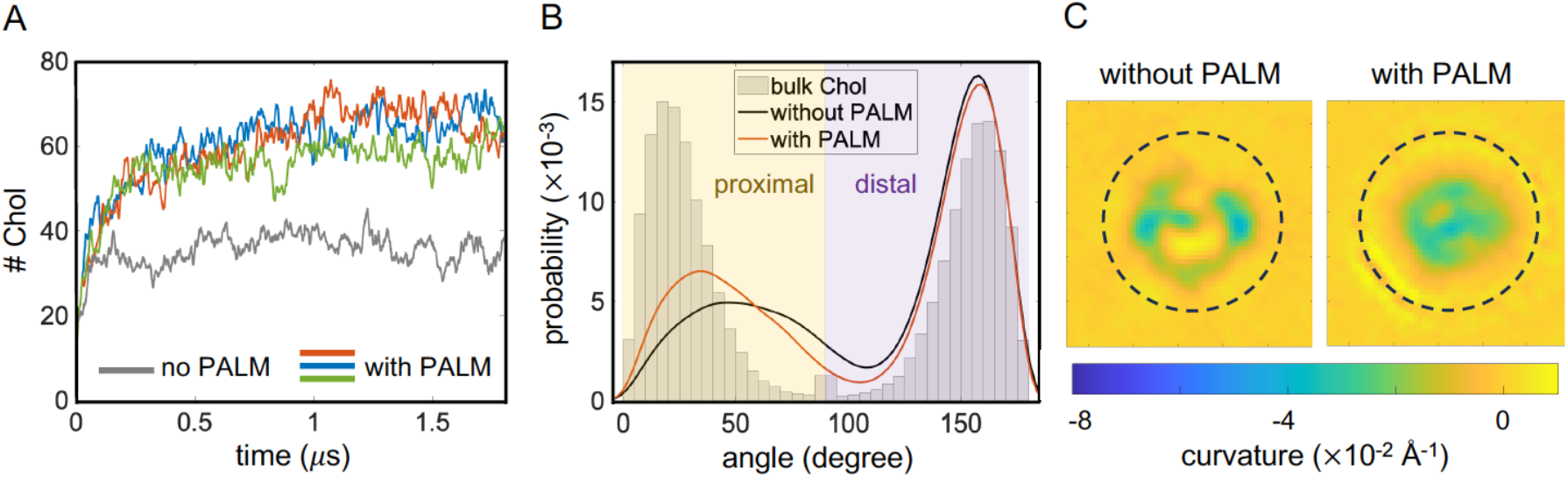
Palmitoylation of caveolin residues affects cholesterol recruitment. (A) Number of Chol molecules in the cytoplasmic leaflet under the complex during the first 1.6 *μ*s of simulation and (B) the equilibrated distribution of their tilt angles with respect to the vector normal to the bilayer surface. Data for the 3 replica trajectories of a POPC+Chol membrane with the palmitoylated CAV1-8S are shown in color; non-palmitoylated are in black. Bars show the distribution of Chol tilt angles in the bulk (i.e. away from the complex). (C) Representative local curvature maps centered on CAV1-8S in simulations with the palmitoylated versus the non-palmitoylated construct. The maps were calculated from the surface of the distal leaflet. Positive curvatures indicate a concave membrane morphology with the protein sitting on top of the curved region. Each map measures 165 × 165 Å and is centered on the complex (shown as dotted circle).

### Association with curved membranes exposes caveolin’s scaffolding domain

Many of caveolin’s reported protein-protein interactions involve residues 81−101 of CAV1, termed the scaffolding domain (SD) (Fig 5A) (65, 66); however, whether SD mediates direct protein binding remains controversial (21, 67-70). This SD region also contains two putative cholesterol-interacting domains known as CRAC (residues 94−101) and CARC (residues 96−103) [reviewed in (21)]. Thus, it is of interest to examine the accessibility of SD residues for binding to lipids or proteins. When CAV1-8S was initially inserted into flat bilayers, many SD residues were buried in the membrane (Fig. 5A, top) due to the hydrophobic nature of this surface. However, several of these SD residues became exposed to the aqueous environment as the membrane around CAV1-8S bent towards the protein (Fig 5A, bottom). This effect can be quantified via residue-specific water-accessible surface area (SASA, Fig. 5B, top), showing that SD residues are generally more solvent-accessible on curved versus flat membranes. This effect was not dependent on lipid composition (Fig. S10).

**Figure 5.**
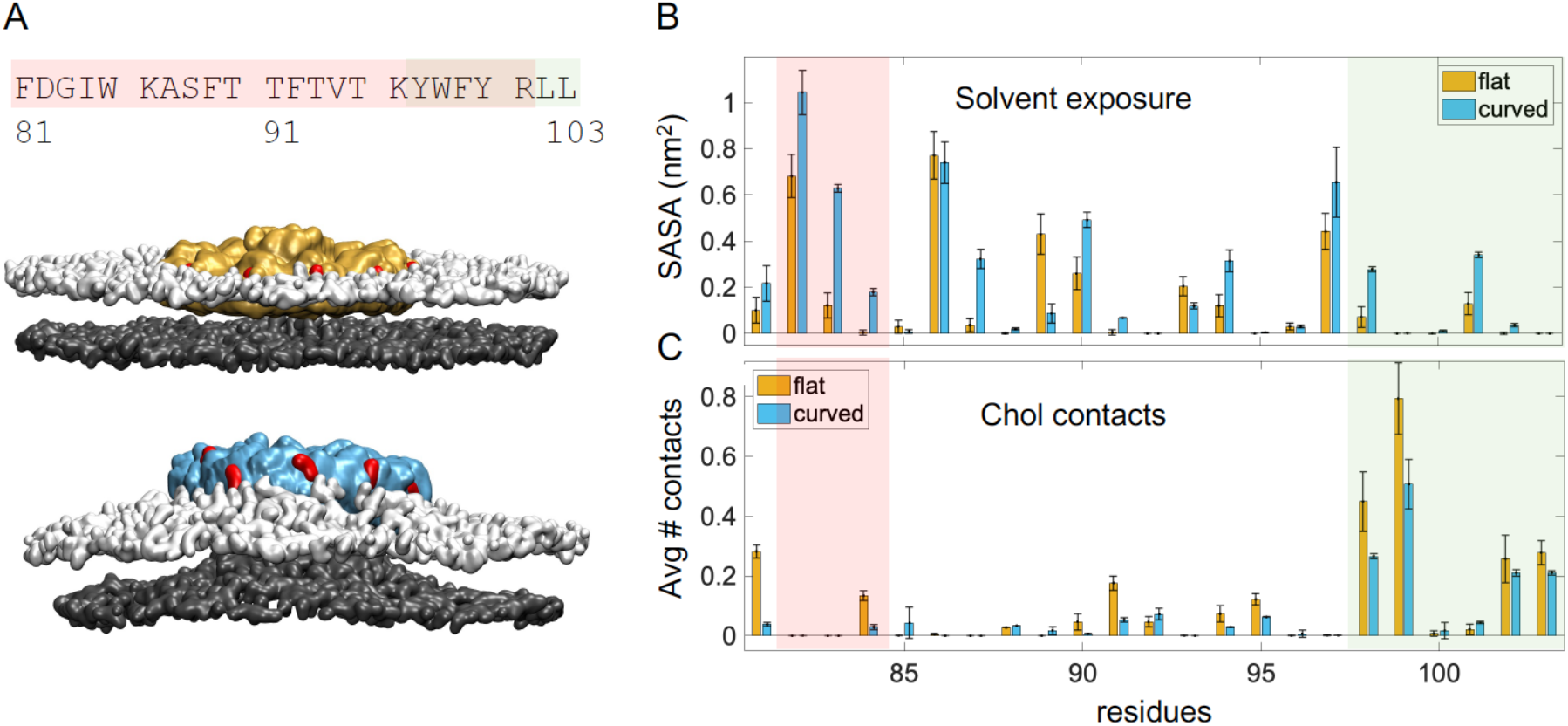
Membrane interactions affect accessibility of caveolin residues to solvent and cholesterol. (A) Residues 81-103 of human CAV1 include the caveolin Scaffolding Domain (SD, 81−101) and a cholesterol-binding motif (CARC, 96−103). Representative snapshots in surface representation of CAV1-8S at the beginning (top, flat) and end (bottom, curved) of the simulation with the asymmetric bilayer (cytosolic leaflet is shown on top). Residues 82−84 of the SD are highlighted in red and the bilayer surface is illustrated by the plane of lipid phosphate groups (cytoplasmic leaflet white, exoplasmic gray). (B) Solvent-accessible surface area (SASA, top) for residues 81−103 and (C) number of contacts with Chol (bottom) averaged over the 11 caveolin protomers from the first 30 ns (flat) and last 1.5 μs (curved) of the simulations. Bars represent the mean and standard deviation across three replica simulations; SD residues 82−84 are shaded in red and CARC residues shaded in green.

Exposure of protein residues to the aqueous environment is expected to reduce their interactions with membrane lipids. To quantify this effect, we measured contacts between SD residues and Chol molecules (Fig. 5B, bottom). Interestingly, CARC (W98, F99, L102 and L103, green shaded region in Fig. 5B) but not CRAC residues showed pronounced direct contacts with cholesterol. In general, these contacts were reduced when CAV1-8S was on a curved membrane surface compared to flat (Fig. 5B, bottom). Thus, the curvature-associated rearrangement of CAV1-8S relative to the membrane has opposing effects on protein-protein versus protein-lipid interactions: on positively curved surfaces, protein-interacting regions become accessible to cytosolic proteins whereas lipid-interacting regions become inaccessible to membrane lipids. These results imply coupling between membrane curvature and intermolecular interactions with caveolin.

## DISCUSSION

The discovery that the human CAV1-8S complex can form a tightly packed flat disc with a large hydrophobic surface challenges long-standing hypotheses about its membrane interactions and prompts new models to explain how caveolin organizes lipids and bends membranes. While the pinwheel structure of the complex has not yet been directly observed *in situ*, two caveolin homologs (from *S. purpuratus* and *S. rosetta*) form similarly shaped oligomeric complexes (71). These observations indicate a conserved biological role of the assembly and highlight the growing need for detailed understanding of its interactions with lipid bilayers. Our computational and experimental findings support a model in which CAV1-8S deeply embeds in membranes by displacing hundreds of lipids from one leaflet of the bilayer (Fig. 1). This arrangement is stable and associated with substantial positive curvature of the membrane towards the caveolin complex, analogous to the curvature of caveolae. We observe substantial lipid sorting by CAV1-8S in both membrane leaflets, most notable in the dramatic recruitment of cholesterol to the proximal leaflet underneath the protein, where it coats the membrane-facing surface of the complex (Figs. 3−4); however, phospholipids are also sorted in the exoplasmic leaflet solvating the protein assembly, with long-chain sphingomyelin preferentially recruited (Fig. 3). These findings may explain caveolins’ reported affinity for cholesterol-rich membranes (21) and their ability to organize lipids in the outer leaflet of the plasma membrane (Fig. 6).

**Figure 6.**
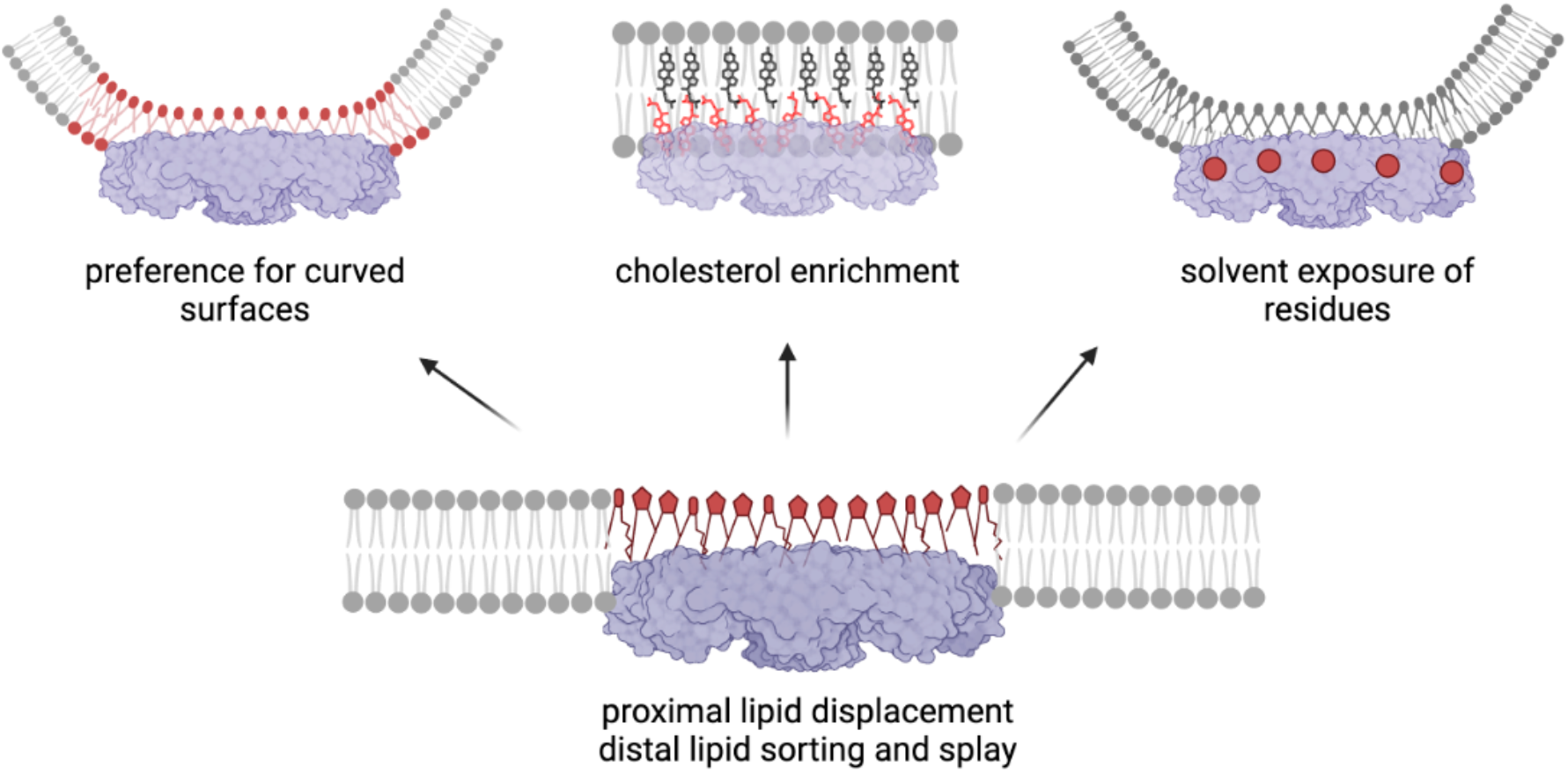
Schematic of the modes of interaction between CAV1-8S and the membrane. The 8S complex embeds deeply into membranes by displacing hundreds of lipids from its proximal leaflet. This arrangement is accompanied by splaying and sorting of the lipids in the distal leaflet, and recruitment of cholesterol molecules underneath the complex into a pure cholesterol monolayer. The complex also has a strong preference for curved membrane regions, which affects the accessibility of putative interaction regions on its surface.

A question raised by this unusual protein-membrane arrangement is how the 8S complex overcomes the large energy barrier that would be required to insert into the membrane during its biogenesis. Current models suggest that CAV1 is synthesized in the endoplasmic reticulum and that the initial membrane insertion of protomers occurs during biosynthesis. In this scenario, assembly of monomers or small oligomers into 8S complexes only occurs after they have trafficked beyond the cis-Golgi complex (72). Thus, it is likely that generation of 8S complexes and accompanying displacement of lipids from the cytoplasmic leaflet in cell membranes occurs gradually as the protein moves through the secretory pathway.

Our observations further suggest that the central β-barrel of CAV1-8S may ‘channel’ cholesterol from the exoplasmic leaflet to the cytoplasm (Fig. 3C), as also suggested by recent metadynamics simulations (58). Our findings come from a simulation that utilized light restraints to maintain the structure of the β-barrel (see Methods). These restraints were necessary because the elastic network typically used in CG simulations to maintain the secondary structure of coarse-grained proteins conserves the structural elements (i.e. α-helices and β-sheets) of individual caveolin monomers, but not structural features arising from their collective assembly (e.g. the β-barrel formed by all 11 C-termini (38), Fig 1A). The restraints applied to maintain the β-barrel did not affect any of the biophysical properties we reported (Fig. S11). Further, we observed spontaneous flipping of Chol molecules from the exoplasmic leaflet into the β-barrel of CAV1-8S in unbiased all-atom simulations (Fig. S8), confirming the ability of the complex to sequester Chol in its central cavity. Longer simulations of these high-resolution models will explore the details of cholesterol movement through the protein complex. Functionally, it is intriguing to speculate how cholesterol sequestration via the complex and/or its presentation via the β-barrel may be related to caveolin’s role in cholesterol trafficking, accessibility, and homeostasis (73, 74).

Best known for their characteristic flask shape, caveolae spend much of their lifecycle in a surface-attached, immobile form. However, it is increasingly recognized that caveolae are not static structures: they function as sensors that flatten and disassemble in response to membrane tension and oxidative stress (15, 20, 26, 31). Thus, the structural and compositional membrane rearrangements induced by CAV1-8S inside and outside of caveolae are critical for the biological functions of caveolin. A key outstanding question is how caveolin bends membranes to form these flask-shaped invaginations. One hypothesis is that CAV1-8S resides on flat regions of the membrane in the absence of cavin-1 or cholesterol, and perhaps even in the flat surfaces of polyhedral caveolae (37). In this model, 8S complexes would be surrounded by patches of free membrane (38), potentially providing space for cavin-1 to associate with lipids and bend the membrane. An alternative scenario proposed in a recent theory (57) and supported by our simulations is that CAV1-8S prefers, and may also induce, highly curved membrane regions, even the vertices of polyhedral structures (46). The underlying physical mechanism of this theory involves an increase in the molecular area of the distal leaflet lipids contacting the protein complex, possibly due to preferential tail-tail interactions in the distal leaflet versus cross-bilayer tail-CAV1-8S interactions. An alternative possibility is that CAV1-8S may displace some of the phospholipids from the distal leaflet. In either case, the increased area/lipid would tend to increase lipid splay and induce membrane curvature, as observed in our CG and atomistic simulations (Fig. 2B and S5). However, other factors likely influence CAV1-8S caveolar biogenesis, including the cholesterol-dependent assembly of caveolins into higher order complexes (75), the N-terminal disordered region of CAV1, cooperative effects of multiple CAV1-8S complexes (58) (Fig. 2D), and other protein-protein interactions. Thus, the full spectrum of molecular interactions that CAV1-8S complexes use to build caveolae remains to be discovered.

Our results also suggest a new structural framework for how caveolin-membrane interactions may regulate the scaffolding function of caveolae. Our observations suggest that the accessibility of several critical SD-domain residues on the outer rim of CAV1-8S is affected by membrane curvature, raising the intriguing possibility that caveolin’s protein and lipid interactions may be regulated by membrane morphology (Fig. 5-6). Thus, the same protein complex may have different sets of interactions inside and outside of caveolae (i.e. in the absence of cavins or when flattened by tension). Finally, the fact that the SD region important for protein-protein interactions contains motifs thought to be responsible for cholesterol binding implies that cholesterol and protein binding events may be mutually exclusive, as also suggested by our simulations.

The recent discovery of the structure of CAV1-8S (38) has provided the context for examining functional caveolin-membrane interactions. Our observations reveal novel modes of protein-lipid interaction and provide mechanistic insight into previous experimental observations of caveolin’s association with specific lipids and membrane morphologies. Further integration of other membrane lipids and proteins into these models is expected to reveal molecular underpinnings of caveolar biology. Further, caveolins appear to belong to a growing class of unusual membranes proteins that bend membranes or sort lipids through large assemblies (including piezos (76), flotillins (77), gasdermins (78), seipins (79)). Exploring the common features of these assemblies may help to reveal novel membrane organizing mechanisms.

## METHODS

### Protein purification

Expression and purification of His-tagged CAV1-mVenus from *E. coli* was performed as previously described (80). CAV1-mVenus was expressed in *E. coli* BL21 using the auto-induction system. Initially, an MDG starter culture was incubated at 37°C with shaking at 250 rpm for 20 hours. Subsequently, the culture was scaled up in auto-inducing ZYM-5052 medium at 25°C and 300 rpm for 24 hours. The *E. coli* cells were then washed with 0.9% NaCl solution and resuspended in a buffer of 200 mM NaCl and 20 mM Tris–HCl, pH 8.0. Bacterial cells were lysed using a French press homogenizer, with 1 mM DTT and PMSF added just before lysis. To eliminate large cellular debris, the lysate was centrifuged at 9000 rpm for 15 minutes at 4°C. Total membranes were collected by centrifugation at 40,000 rpm (Ti-45 rotor, Beckman Coulter) for 1 hour at 4°C. The membrane pellets were then homogenized using a Dounce tissue grinder in a buffer of 200 mM NaCl, 20 mM Tris–HCl (pH 8.0), and 1 mM DTT. To extract caveolin proteins from the membranes, a 10% n-Dodecyl-β-D-maltoside (C12M) (Anatrace) stock solution was added to the membrane homogenate to a final concentration of 2%, and the mixture was gently agitated for 2 hours at 4°C. Insoluble material was separated by centrifugation at 42,000 rpm (Ti-50.2 rotor) for 35 minutes at 4°C, and the supernatant was subjected to nickel Sepharose affinity purification. The eluate containing caveolin was concentrated and further purified by size-exclusion chromatography using a Superose 6 Increase 10/300 GL column (GE Healthcare) in a buffer containing 200 mM NaCl, 20 mM Tris–HCl (pH 8.0), 1 mM DTT, and 0.05% C12M. Purified Syntaxin1a and SNAP25 were kind gifts of Katelyn Kraichely (Lukas Tamm lab, UVA) (81).

### Film balance experiments

Experiments were carried out on a DeltaPi-4x film balance setup (Kibron, Finland), which is equipped with four identical Teflon troughs for parallel measurements. Each trough has a fixed, circular area of 4.075 cm^2^. For measurements of surface pressures with the Wilhelmy method, a metal DyneProbe is suspended from a microbalance above each trough. The subphase consisted of 2.1 mL of 200 mM NaCl and 20 mM Tris-HCl (adjusted to pH 8.0 with NaOH). Measurements were performed at 20°C with the subphase constantly being stirred with a magnet stirrer. During experiments, a hood with wet tissues on the inside was placed over the setup to maintain a humid atmosphere thus limiting water evaporation from the troughs. Before each experiment, the trough was cleaned with 6 M guanidinium chloride and Hellmanex III-solution (Hellma, Germany) and then thoroughly rinsed with ultrapure water (Millipore, Germany). The trough was filled with buffer, the microbalance was calibrated, and the probe placed at the surface. At this point, the surface pressure value was set to zero and a pressure baseline was recorded for a minimum of 30 min. For insertion measurements at the liquid-air interface, different amounts of protein or buffer (200 mM NaCl, 20 mM Tris-HCl pH 8.0, 1 mM DTT, 0.05% C12M) were injected with a Hamilton syringe under the surface, through the trough’s side port. Purified CAV1-8S was either used directly or after concentrating with an Amicon filter concentrator with a molecular weight cutoff of 100 kDa (Millipore, Germany). In the latter case, the buffer for the control was treated in the same way. Changes in surface pressure were continuously recorded for several hours after injection of protein or buffer control. For lipid monolayer insertion experiments, a lipid film was spread on the buffer surface from a 0.1 mM solution of pure lipids (Avanti) in chloroform/methanol (2:1 volume ratio) using a Hamilton syringe. After the rapid evaporation of the organic solvent, the film was allowed to stabilize for approximately 5 h before protein or buffer were injected into the subphase. Changes in surface pressure were continuously recorded for several hours after sample injection.

### Preparation of liposome suspensions

Large unilamellar vesicles (LUVs) were prepared by extrusion (54). POPC (Avanti) and Texas-Red DHPE (Invitrogen) were mixed in an amber vial (2 mL, Agilent) to yield a mixed lipid solution with a molar ratio of 99.05 POPC:0.05 Texas Red-DHPE. Excess solvent (chloroform) was evaporated under gentle nitrogen stream to form a damp lipid film, and trace chloroform was removed by drying for 2 h under house-vacuum. After drying, the mass of the dried lipid film was determined. LUV buffer (360 mM sucrose, 25 mM NaCl, and 25 mM Tris pH 7.4) was added to resuspend the lipid film. This suspension was flash-frozen by submerging sealed vials into liquid nitrogen and then rapidly thawed by transferring to 50°C water bath; repeated five times. Then, using an extruder (Avanti, 610000), the suspension was passed 23 times through a 100 nm polycarbonate filter (Whatman, 19 mm).

### HPaCD-mediated reconstitution of CAV1-8S into LUVs

Mixtures containing LUVs, HPaCD, and CAV1-8S were prepared with lipid concentrations fixed at 0.5 mM and 19 nM CAV1-8S, 31 mM NaCl, 0.002% C12 M (m/v%), 7.4 mM sodium cholate, and 25 mM Tris pH 7.4 in the external LUV solution. To maintain similar osmotic pressure across the LUVs in each mixture, sucrose was used as a balancing osmolyte: in each mixture, it was ensured that the combined concentrations of HPαCD and sucrose in the external solution combined to 360 mM, matching the sucrose concentration encapsulated in the LUVs. Mixtures were prepared by first adding LUVs then CAV1-8S (in rapid succession) into the aqueous solutions while stirring with a magnetic stir bar for 20 min at 23°C. After the 20 min reaction, samples were dialyzed against LUV buffer supplemented with 1−2 g of Bio-Beads SM-2 resin (Bio-Rad) overnight at 4°C to remove detergent using 1 kDa MWCO Tube-O-DIALYZER (G-Biosciences).

### Co-flotation assay

Liposome samples collected from dialysis were mixed with an equal volume of 1.75 M sucrose (25 mM NaCl, 25 mM Tris pH 7.4, bottom layer). 760 mM sucrose (in same buffer) was layered on top (middle layer) followed by LUV buffer (top layer). This sucrose layers were spun at 185,000x*g* for 1 h at 4°C (Ti-70.1 rotor, Beckman Coulter). After centrifugation, samples were fractionated from the top in 100 μL aliquots into a FLUOTRAC 96-well plate (Greiner Bio-One). To measure relative protein and lipid amount per fraction, excitation/emission was set to 505/595 nm for Cav1-mVenus and 595/615 nm for Texas-Red DHPE.

### Monitoring solubilization of LUVs by HPaCD

90° static light scattering of 380 nm light was used to observe vesicle solubilization by HPaCD (54). Measurements were performed in the absence of CAV1-8S but with otherwise identical concentrations as used for HPaCD-mediated reconstitution. A Horiba Fluorolog operated in “synchronous scan” mode was used to collect scattering from samples in quartz microcuvettes (Starna Cells, Inc., 10 mm pathlength).

### Dynamic Light Scattering (DLS) of LUV/HPaCD/CAV1-8S mixtures

A Malvern Zetasizer nano series ZS90 (633 nm laser) was used to determine the size-distribution of particles in LUV/HPaCD/CAV1-8S reconstitution mixtures. The particle size-distribution was determined from the fluctuations in intensity of scattered light assuming viscosity of 0.8872 cP for the aqueous dispersant and spherical particles.

### Design and construction of simulation systems

We simulated five different coarse-grained CAV1-8S-membrane systems: bilayers containing 1, 2, 6 or 9 caveolin complexes without acylation, and bilayers with 1 fully palmitoylated-caveolin complex (Table S2).

#### Protein

The CAV1-8S structure was taken from the recently deposited cryoEM data of the human Caveolin-1 8S complex (PDB 7SC0). Each of the 11 protomers contained residues 49 through 177 with neutral terminal caps; the unstructured N-terminal 48 amino acids are omitted from our model, as they were not included in the cryoEM structure. The palmitoylated model was first constructed in all-atom representation with CHARMM-GUI PDB Reader, then it was converted to coarse-grained representation with the *martinize* script.

#### Bilayers

The protein constructs were incorporated into four types of bilayers: (1) ***POPC***, pure POPC, (2) ***POPC+Chol***, POPC with 30 mol% Chol, (3) ***inner***, symmetric mixture of 1-palmitoyl-2-arachidonoyl-sn-glycero-3-phospho-L-serine (PAPS), 1-palmitoyl-2-linoleoyl-sn-glycero-3-phosphocholine (PLPC), 1-palmitoyl-2-docosahexaenoyl-sn-glycero-3-phosphoethanolamine (PDPE) and cholesterol (Chol) with molar ratios of PAPS/PLPC/PDPE/Chol 26/18/16/40, and (4) ***asym***, an asymmetric bilayer with CAV1-8S-proximal leaflet with the *inner* lipid composition and CAV1-8S-distal leaflet made of N-palmitoyl-D-erythro-sphingosylphosphorylcholine (16:0 SM or PSM), N-nervonoyl-D-erythro-sphingosylphosphorylcholine (24:1 SM or NSM), PLPC and Chol with molar ratios of PSM/NSM/PLPC/Chol 16/14/30/40.

The following coarse-grained MARTINI lipids were used in the simulations: PAPS (for PAPS), PRPE (for PDPE), PIPC (for PLPC), DPSM (for PSM) and PNSM (for NSM). All Chol-containing bilayers were built and equilibrated with the older 2.2 version of CHOL, however the molecule was replaced with the newer CHOL (82) model prior to the production runs for most systems. The only exceptions were 1 of the replicas of the *inner* bilayer with a single CAV1-8S, 1 of the replicas of the *inner* bilayer with two 8S complexes and all bilayers with 6 or 9 complexes. No measurable differences were observed between the two CHOL models.

#### Protein-bilayer systems

The single-CAV1-8S systems with the non-palmitoylated construct were built with the Martini Maker tool in CHARMM-GUI using the Positioning of Proteins in Membranes (PPM) web server’s prediction to guide the placement of the complex inside one of the bilayer leaflets. For initial construction, the CAV1-8S-proximal leaflet contained 250 lipids (including cholesterol) fewer than the distal leaflet. The bilayer with the palmitoylated 8S complex was built with the *insane* tool and its proximal leaflet had 254 lipids fewer than the distal.

The systems with 2, 6 and 9 CAV1-8S complexes were assembled manually from copies of single-CAV1-8S bilayers using the Visual Molecular Dynamics (VMD) software (Table S1). The initial lateral distance between the centers of mass of neighboring complexes were: 45 nm (for 2-CAV1-8S), 17 nm (for 6-CAV1-8S) and 22 nm (for 9-CAV1-8S). Topology files were generated with in-house MATLAB scripts. The systems were solvated and ionized with Gromacs.

All-atom systems of CAV1-8S in a POPC or POPC/Chol 70/30 bilayer were constructed with CHARMM-GUI’s membrane builder tool. They were about 300 × 300 × 160 Å in size and contained 150 mM KCl. Each protein monomer was modeled with a neutral N-terminus and a standard charged C-terminus.

### Simulation software and parameters

All coarse-grained simulations were run with Gromacs (83), the Martini v2.2 force field for proteins and the Martini v2.0 (from 201506) force field for ions and lipids (except for CHOL whose model and parameters were updated to a newer model (82) for the production runs, as explained above). The parameters for the palmitoyl chains were taken from (84). Equilibration was performed following CHARMM-GUI’s 6-step protocol followed by a short 30 ns simulation and a longer production run (see Table S1). The simulation parameters included a time step of 20 fs, a VdW cutoff of 1.1 nm applied with a potential shift, reaction-field electrostatics with a cutoff of 1.1 nm, a neighbor list update every 20 steps and lincs_iter and lincs_order parameters set to 1 and 4 respectively. Temperature was maintained at 37 °C (310.15 K) by velocity rescaling and the system was coupled to a Parrinello-Rahman barostat for semi-isotropic pressure control.

An additional coarse-grained simulation of CAV1-8S in the POPC+Chol bilayer was run while restraining the structure of the beta-barrel. This was achieved by applying harmonic potentials to all pairwise distances between the backbone beads of residues N173 and E177 in all 11 caveolin protomers. The distances were restrained to those from the cryo-EM structure using the distance geometry in Gromacs in all 3 dimensions with a pulling rate of 0.0 nm/ps and a force constant of 1000 kJ mol^-1^ nm^-1^.

All-atom simulations were performed with Gromacs and the Charmm36 force field (85, 86). The simulations were run with a 2-fs timestep and bonds with hydrogens were constrained with the Lincs algorithm. Non-bonded interactions were modeled with the Verlet cut-off scheme, a 10−12Å Lennard-Jones potential with a force-switching function and fast smooth Particle-Mesh Ewald electrostatics with 1.2 nm cutoff. Temperature (310.15 K) and pressure (1 bar in z) were controlled with the v-rescale and C-rescale algorithms, respectively. Three 1-μs long replica simulations were run for each system, starting from the same structures and regenerating the initial velocities.

### Analysis of simulation trajectories

All analysis was performed with either Gromacs or in-house Tcl, bash and MATLAB scripts. The VMD software was used for visualization and structure rendering. Unless otherwise noted, quantitative analyses were performed on the final 1.5 μs (for coarse-grained) or 300 ns (for all-atom) of trajectories, where the RMSD was equilibrated (Fig. S12).

#### Lipid composition

Lipids under CAV1-8S versus bulk were counted based on the location of their headgroups: they were considered under the protein complex if the (x,y) coordinates of their PO4 beads were within 2 Å of the (x,y) coordinates of any protein bead. Assignment to cytoplasmic versus exoplasmic leaflet was determined based on the orientation of the vector connecting the lipid headgroup PO4 bead to the chain C3B bead. Chol leaflet assignment was determined by its tilt with respect to local vectors normal to the surface under the protein, as described below, or with respect to the bilayer normal (i.e. the z dimension of the simulation box) in the bulk.

#### Cholesterol tilt

Chol tilt was calculated from the orientation of the director vector connecting its headgroup ROH bead and a bead in its rigid ring body (C5). In the bulk, Chol tilt was obtained from the angle between this vector and the bilayer normal (z dimension of the simulation box). Due to the high curvature associated with CAV1-8S, under the protein Chol tilt was calculated from the angle between its director vector and local bilayer normal, calculated as the vectors normal to the surface interpolated from the PO4 bead positions of the lipids in the leaflet distal to CAV1-8S.

#### Curvature and chain splay maps

A surface of the distal membrane leaflet was created in every frame of the trajectory by interpolating the 3D positions of the lipids’ PO4 (or P for the all-atom systems) beads after aligning the protein backbone atoms (to keep orientation consistent). The curvature was calculated with a lateral resolution of 3 Å on a frame-by-frame basis with MATLAB’s *surfature* function and averaged over the entire trajectory. From the same aligned trajectory, the chain splay of the lipids in the distal leaflet was analyzed by calculating the angle between the two vectors defining the chain orientations (i.e. vectors connecting the terminal bead to the corresponding glycerol backbone bead; in the all-atom systems these corresponded to atoms C31 and C316, and C21 and C218). Both the curvature and chain splay were mapped onto the same 2D grid for visualization.

#### Membrane deformation in bilayers with two CAV1-8S complexes

To monitor the evolution of the large deformations in the symmetric bilayer (with ***inner*** lipid composition) having one CAV1-8S complex embedded in each leaflet, we calculated the maximal vertical distances (i.e. only in the z dimension) between either (1) lipid headgroups or (2) the two oppositely situated complexes in each frame. For the former, we used the distance between the minimum and maximum z coordinates across all lipid PO4 beads; for the latter, the distance between the maximum z coordinate across all beads of CAV1-8S in the top leaflet and the minimum z coordinate across all beads of CAV1-8S in the bottom leaflet. Both measures showed the same trend and quantitatively captured the gradual formation of large curvature in the trajectories.

#### Solvent exposure of residues

To quantify the exposure of caveolin residues to solvent, we calculated the solvent-accessible surface area (SASA) of each residue in every trajectory using the *sasa* tool in Gromacs with a probe radius of 2.6 Å and VdW radii consistent with the coarse-grained representation of the molecules: 2.64 Å for protein backbone and lipid carbon beads, 2.3 Å for a protein side chain bead, 3.48 Å for the choline headgroup and PO4 beads, and 2.41 Å for the ROH bead. ΔSASA was derived by comparing the last 1.5 μs of the trajectory to the first 30 ns of the short production run (when all systems were essentially flat).

#### Cholesterol contacts with CAV1-8S

Contacts between caveolin residues and cholesterol were defined as proteins residues being within 5 Å of any cholesterol bead averaged over trajectory frames and the 11 caveolin protomers. We repeated the calculation by restricting it to only residues in contact with Chol’s ROH headgroup bead and observed no significant differences.

## DATA, MATERIALS AND SOFTWARE AVAILABILITY

All simulation trajectories, analysis scripts, and experimental data will be made available on Zenodo and GitHub.

## ACKNOWLEDGMENTS

We thank Richard Pastor, Helgi Ingolfsson, Gregory A. Voth and Korbinian Liebl for helpful discussions. Funding for this work was provided by the National Institutes of Health (NIH)/National Institute of General Medical Sciences (GM134949 to IL, 1F32GM134704-01 to MD, R01HL168258 to AKK, R01GM138444 to PMK) and Deutsche Forschungsgemeinschaft (436494874-RTG2670 to SN/KB). Computational resources were provided by the National Academic Infrastructure for Supercomputing in Sweden (NAISS). In addition, this work used the Delta supercomputer in the National Center for Supercomputing Applications (NCSA) at University of Illinois Urbana-Champaign through allocations MCB180168 and BIO240103 from the Advanced Cyberinfrastructure Coordination Ecosystem: Services & Support (ACCESS) program, which is supported by National Science Foundation grants #2138259, #2138286, #2138307, #2137603, and #2138296. Some of the figures were created with BioRender.

## AUTHOR CONTRIBUTIONS

MD, IL, AKK, KB, TRR, HIC, and KRL designed research; MD, PK, SD, JE, SN, BH, TRR, HIC performed research; MD, SS, PK, SD, JE analyzed data; MD, IL, AKK, KB and KRL wrote the paper.

