## Supplemental Material for "Caveolin assemblies displace one bilayer leaflet to organize and bend membranes"

The document includes 2 tables and 12 figures.

**Table S1. Monolayer maximum insertion pressure (MIP) measured from Langmuir film experiments.** Shown are the lipid composition of the monolayer and the MIP with 95% confidence intervals obtained from the data in Figs. 1D and S2C–E.

| Monolayer composition | protein | MIP / mN/m | 95% CI / mN/m |
|---|---|---|---|
| POPC | CAV1-8S | 33.0 | 31.4 – 34.7 |
| POPC/Chol | | 37.8 | 31.6 – 43.9 |
| POPC/POPE/POPS | | 41.1 | 39.2 – 43.0 |
| POPC/POPE/POPS | Sar1-ΔN | 24.6 | 22.8 – 26.4 |

**Table S2. Simulated membranes containing Cav8S.** Shown are the number of protein complexes in the bilayer; the membrane lipid composition; the number of total lipids (including Chol) in each leaflet at the time of construction; the box dimensions at the time of construction; the total number of replica simulations and how many of those were used for analysis; and the total length of the production runs used for analysis. All systems had 150 mM salt concentration (Na and Cl ions) unless otherwise noted, and were simulated at a temperature of 37 °C (310.15 K).

| # Cav8S | lipid composition | # total lipids at t=0 | | Box size at t=0 [nm] | # replicas | | production run [$\mu$s] |
|---|---|---|---|---|---|---|---|
| | | proximal | distal | | total | analysis | |
| 1 | POPC* | 3,250 | 3,500 | 49×49×22 | 3 | 1 | 8 |
| | POPC/Chol 70/30 | 3,250 | 3,500 | 45×45×26 | 3 | 1 | 22 |
| | Inner[1] | 3,250 | 3,500 | 45×45×26 | 4 | 3 | 7, 10, 10 |
| | Asym[2] | 3,250 | 3,500 | 45×45×26 | 6 | 3 | 6.6, 8, 10 |
| 1-PALM[3] | POPC/Chol 70/30 | 3,280 | 3,534 | 42×42×35 | 3 | 3 | 10, 10, 10 |
| 1-Beta[4] | POPC/Chol 70/30 | 3,250 | 3,500 | 45×45×26 | 1 | 1 | 11.6 |
| 2 | Inner[1] | 6,750 | 6,750 | 92×46×24 | 3 | 3 | 2.8, 3.2, 3.8 |
| 6 | Inner[1] | 13,854 | 15,268 | 78×116×26 | 3 | 3 | 23.5, 35.7, 38.2 |
| 9 | Inner[1] | 27,850 | 30,185 | 158×114×30 | 3 | 3 | 12.7, 17.2, 16.8 |
| 0 | Inner[1] | 29,636 | 29,636 | 158×114×30 | 4 | 4 | 5.8,5.6,5.6,5.6 |
| 0 | Inner[1] | 6,269 | 6,269 | 83×42×35 | 1 | 1 | 3 |
| 1 | POPC AA[5] | 1,123 | 1,318 | 30×30×16 | 3 | 3 | 1 |
| 1 | POPC/Chol 70/30 AA[5] | 1,290 | 1,510 | 30×30×16 | 3 | 3 | 1 |

[1]symmetric membrane mixture of PAPS/PLPC/PDPE/Chol 26/18/16/40 mol%.
[2]asymmetric membrane with proximal leaflet of PAPS/PLPC/PDPE/Chol 26/18/16/40 mol% and distal leaflet of PSM/NSM/PLPC/Chol 16/14/30/40 mol%.
[3]palmitoylated CAV1-8S with 33 palmitoyl chains (3 per protomer) constructed with the insane.py tool
[4]simulation in which the structure of the central $\beta$-barrel was restrained during the trajectory
*this system was electro-neutralized with sodium ions but did not have any added salt
[5]all-atom simulations of membrane-embedded CAV1-8S in the presence of 150 mM KCl

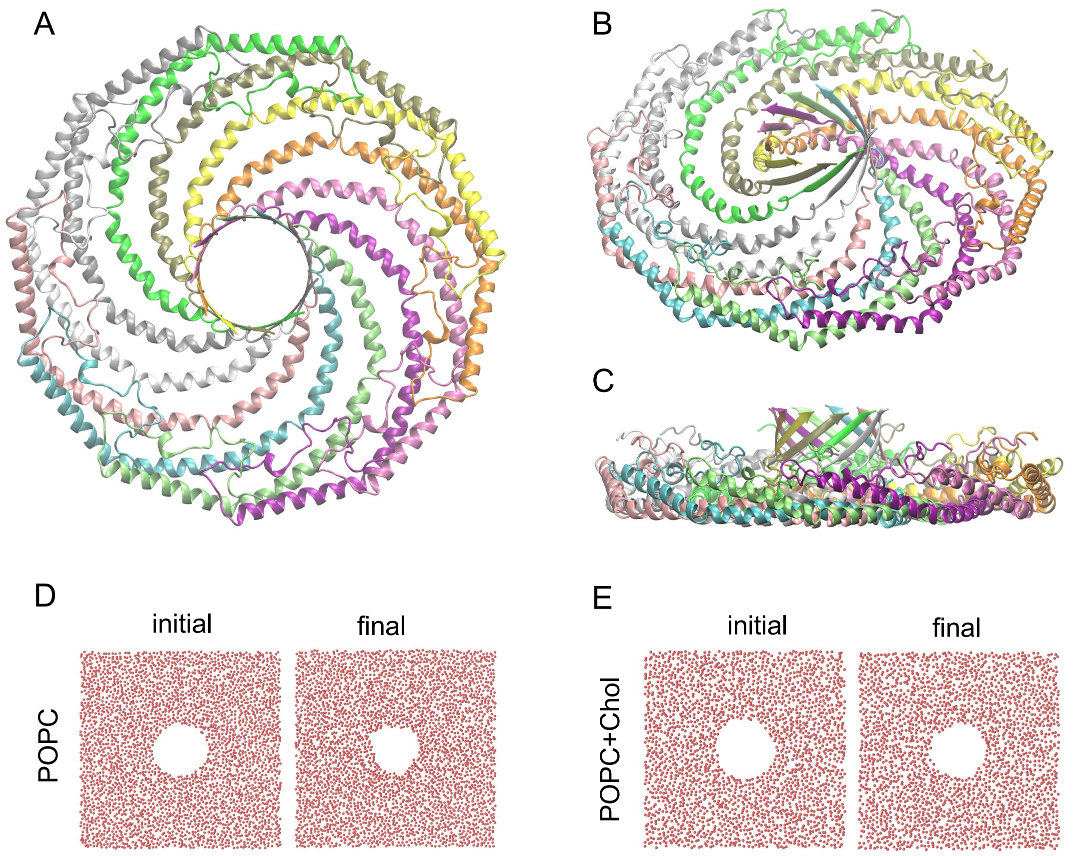


**Figure S1. Structure and membrane interaction of CAV1-8S.** (A−C) Spiral arrangement of CAV1 monomers in the 8S complex. Each one of the 11 monomers is colored in a different color. Panels show different views: top (A), tilted (B) and side (C). (D−E) Phospholipid distribution in the CAV1-8S-proximal leaflet (viewed from above the leaflet) at the beginning (left) and end (right) of the POPC (D) and POPC/Chol 70/30 (E) lipid bilayers. Red dots represent positions of phospholipid phosphate groups in the bilayer leaflet proximal to the protein complex.

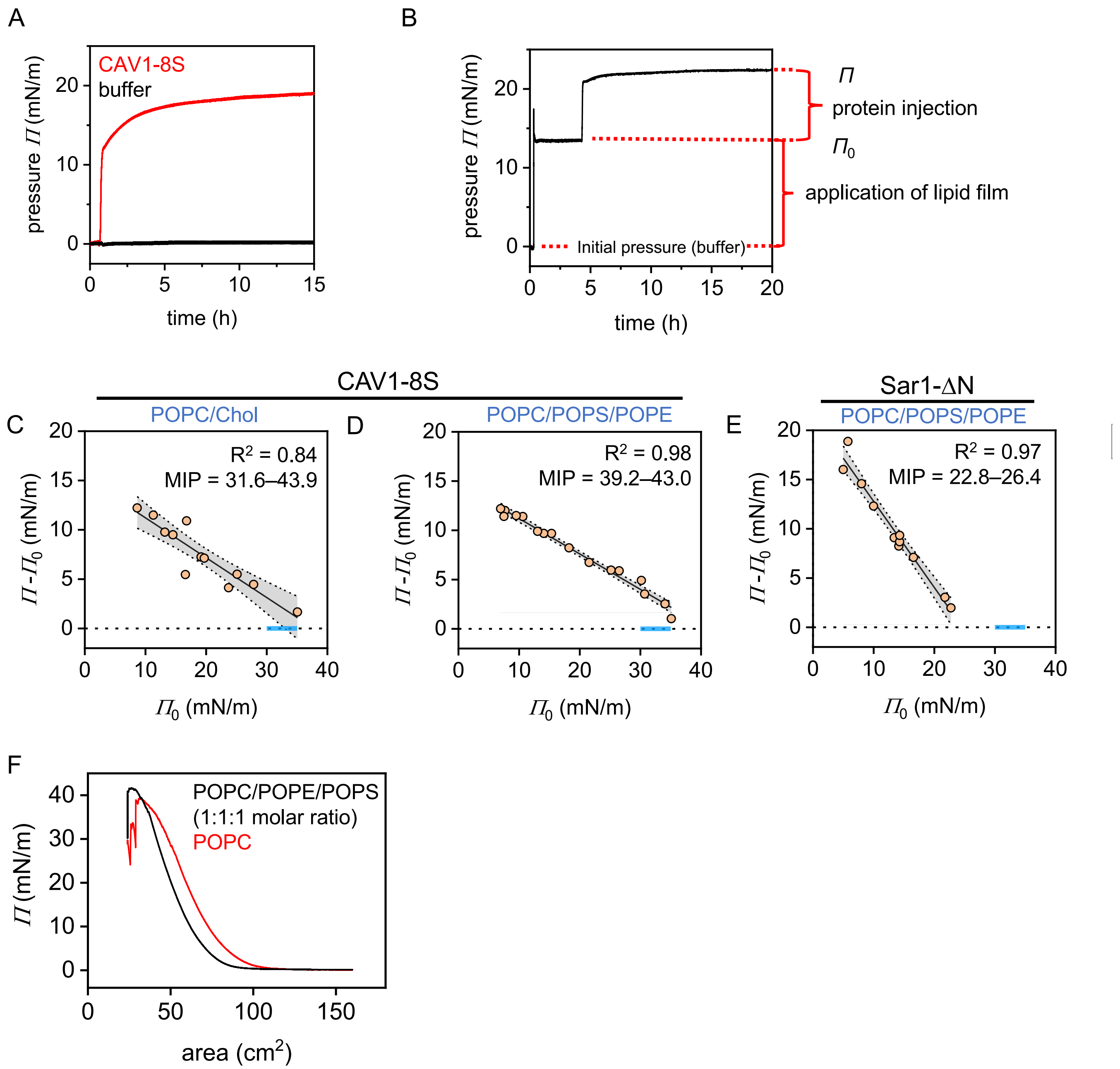


**Figure S2. CAV1-8S incorporates into tightly packed lipid monolayers.** (A) Addition of CAV1-8S to the aqueous subphase in the Langmuir trough results in rapid increase in surface pressure, which equilibrates at an elevated value indicating high surface activity of the protein complex. (B) Application of a lipid film to the water/air interface produces an elevated surface pressure $\Pi_0$ upon evaporation of the organic solvent. Subsequent injection of protein increases the pressure further to $\Pi$. The difference $\Delta\Pi = \Pi - \Pi_0$ as a function of $\Pi_0$ is used to characterize protein binding to a lipid monolayer. (C) $\Delta\Pi$ as a function of $\Pi_0$ for insertion of CAV1-8S into a POPC/Chol (3:1 molar ratio) monolayer. (D) Same as (C) but for a monolayer composed of 1:1:1 POPC/POPS/POPE. (F) $\Delta\Pi$ as a function of $\Pi_0$ for insertion of Sar1-ΔN (Sar1 lacking its membrane-inserting N-terminus) into a 1:1:1 POPC/POPS/POPE monolayer. The blue bar in (C), (D) and (E) indicates the equivalence pressure (EQP, between 30–35 mN/m), at which the lipid packing in the monolayer is comparable to that in a bilayer. (F) Pressure-area isotherms of POPC and 1:1:1 POPC/POPS/POPE monolayers.

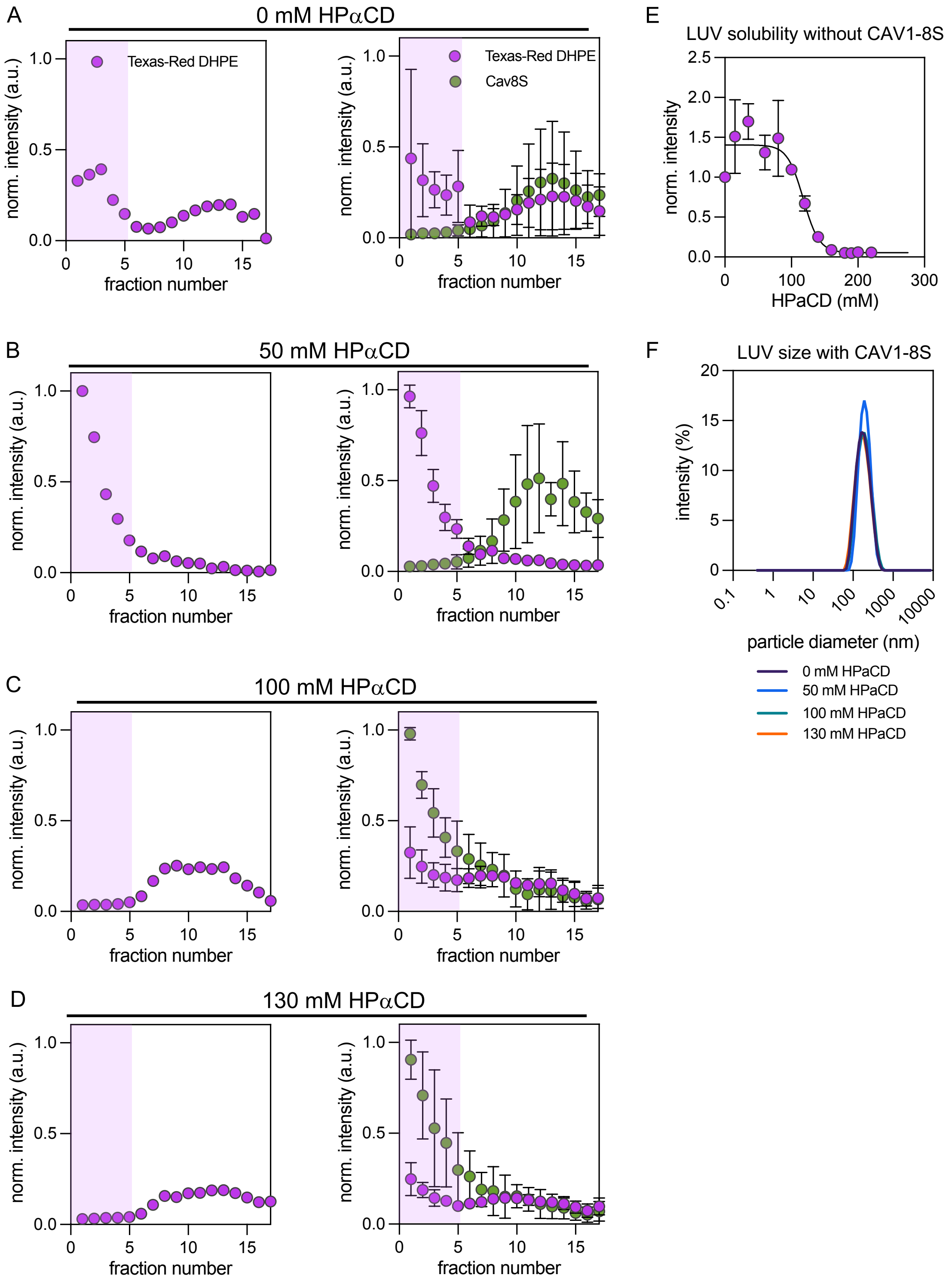


**Figure S3. Insertion of CAV1-8S into model liposomes.** (A-D) Co-flotation assays of control POPC vesicles with liposomes only or with CAV18S-mVenus at increasing HPaCD concentrations. Top, liposome-containing fractions (1-5) are highlighted in purple. N=1 for control liposomes, N=3 for all samples with CAV1-8S, errors bars are standard errors of the means (SEMs). (E) Light scattering intensity reports on intact LUVs and their disruption by HPaCD in the absence of CAV1-8S. Intensities are normalized to 0 mM HPaCD. N=3, error bars are SD. (F) Size distribution from DLS of fraction 1 from co-floatation assays in A-D.

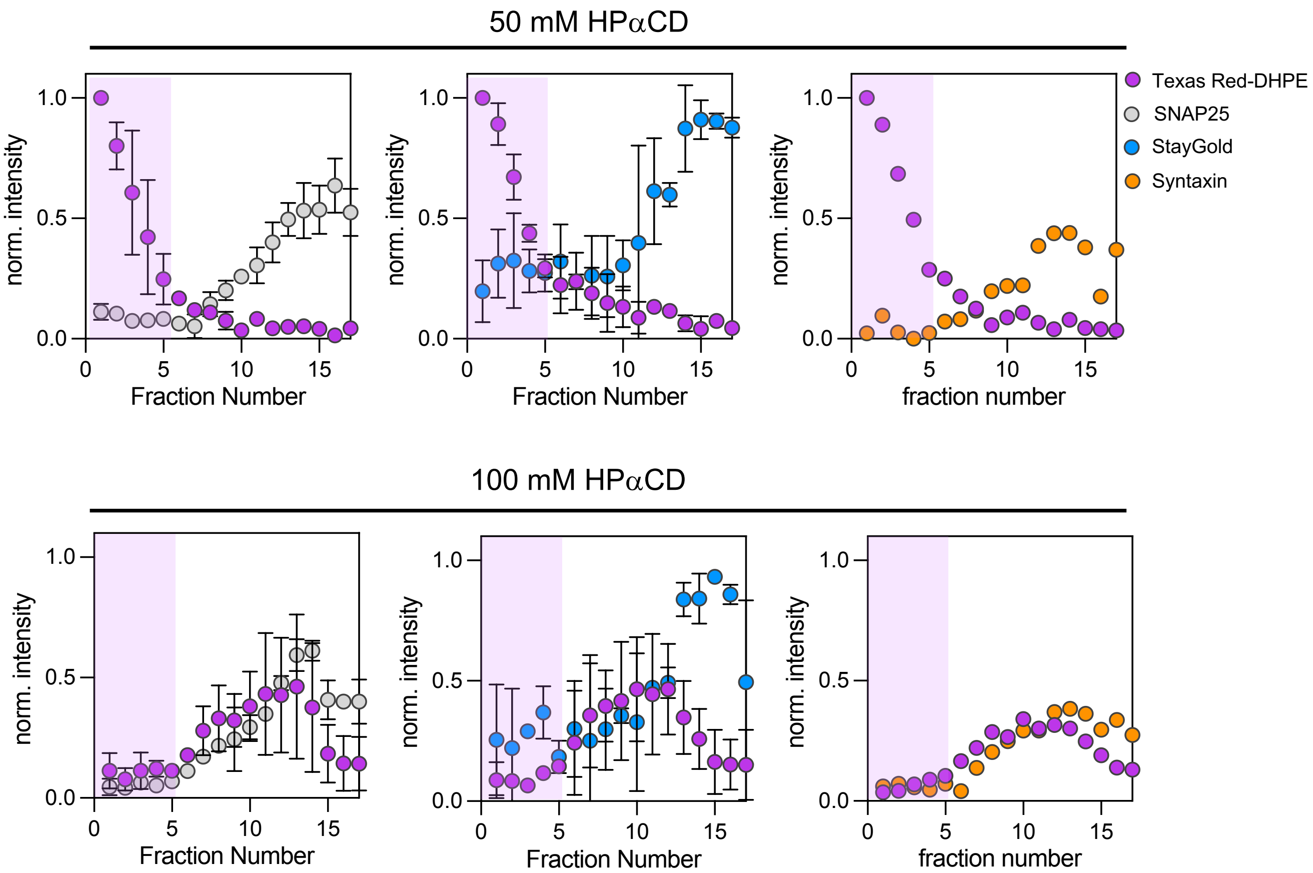


**Figure S4. Effects of cyclodextrin on binding of non-caveolin proteins to model liposomes.** Co-flotation assays of POPC liposomes with recombinantly purified SNAP25-AF647 (grey), StayGold-GFP (blue), and Syntaxin1a-AF647 (orange) at 50 mM (top) and 100 mM (bottom) HPaCD. HPaCD had no effect on incorporation of any of these control proteins.

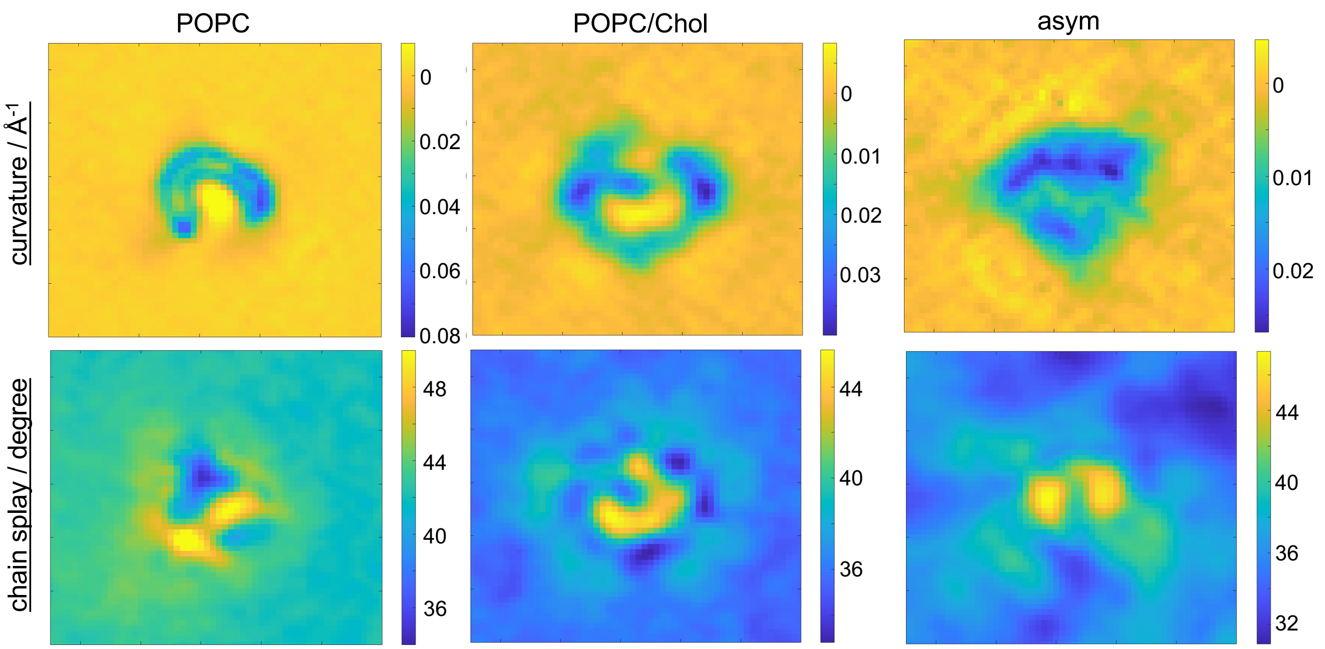


**Figure S5. Curvature and chain splay in coarse-grained CAV1-8S-membrane simulations.** Maps of the curvature (top) and lipid chain splay (bottom) of the CAV1-8S-distal leaflet in the POPC, POPC/Chol and asymmetric bilayers. Each map is 165 × 165 Å and is centered on the protein complex. More positive curvatures indicate monolayer bending towards CAV1-8S (see Fig. 2B) while larger positive splay angles correspond to lipids with more splayed-out chains and larger areas per lipid.

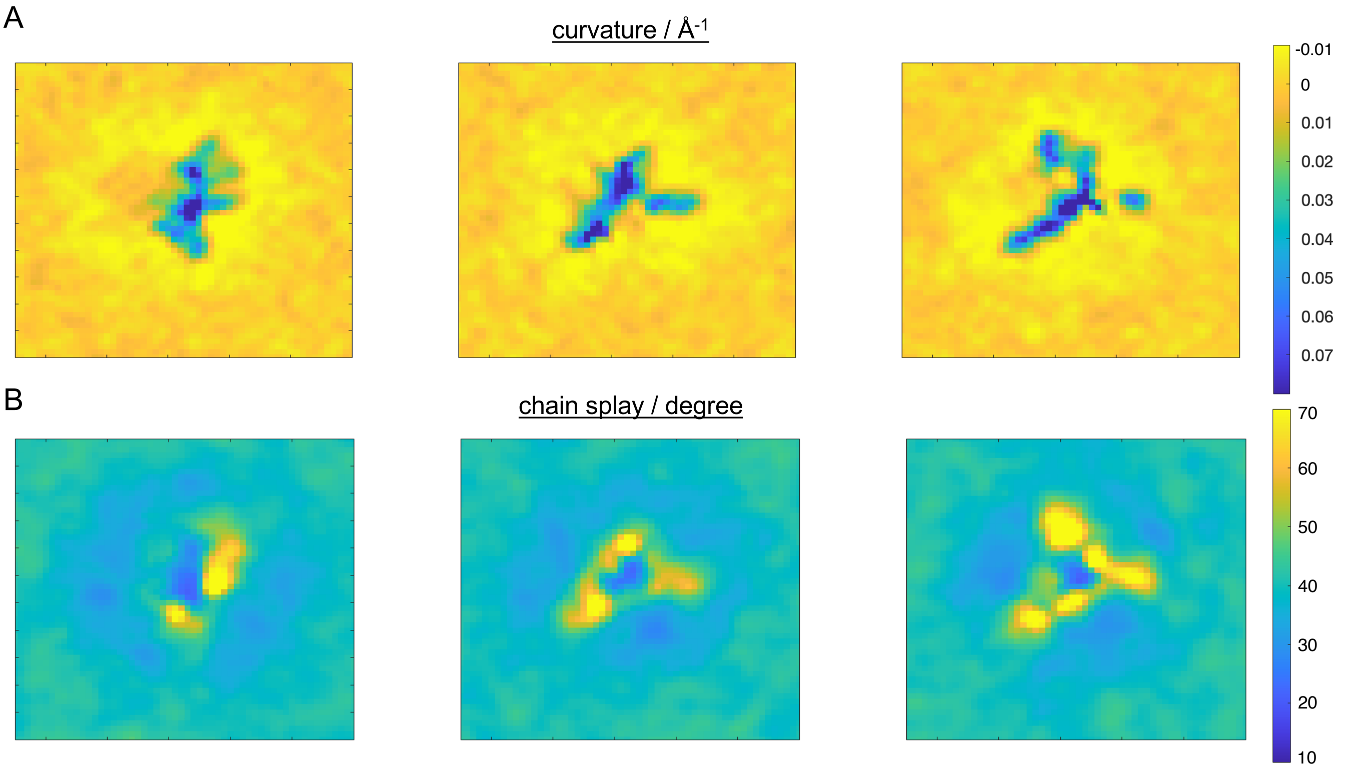


**Figure S6. Curvature and chain splay in all-atom CAV1-8S-membrane simulations.** Maps of the curvature (A) and lipid chain splay (B) of the CAV1-8S-distal leaflet in three all-atom replica simulations of the complex embedded in a POPC membrane. Each map is 165 × 165 Å and is centered on the protein complex as in Figure S5.

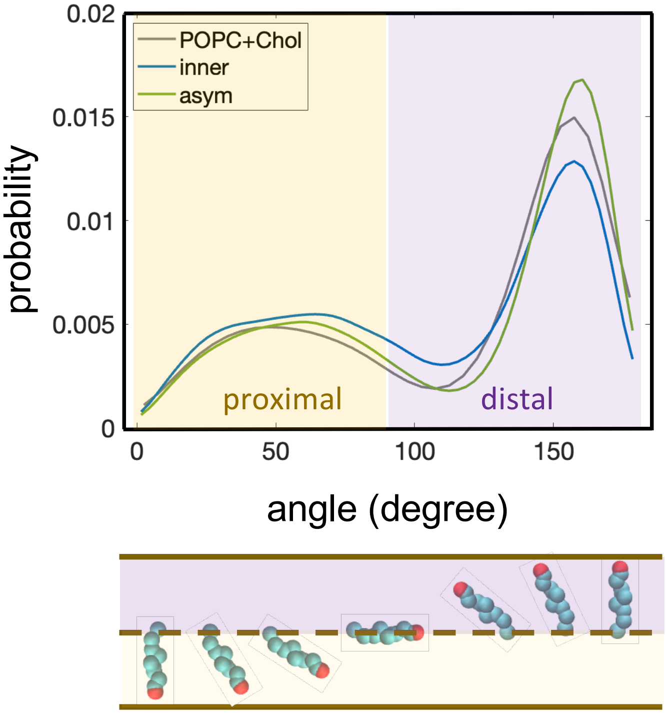


**Figure S7. Orientation of cholesterol molecules in coarse-grained CAV1-8S-membrane simulations.** Equilibrated distribution of Chol tilt angles with respect to the bilayer normal in the three simulated bilayers, POPC+Chol, inner and asym. The bilayer normal was calculated locally as explained in Methods to avoid any bias due to the curvature in the systems. The data shown for the inner and asym bilayers is the average of 3 replica simulations. Bottom schematic illustrates the orientation of a Chol molecule corresponding to the respective tilt angles enumerated on the $x$ axis.

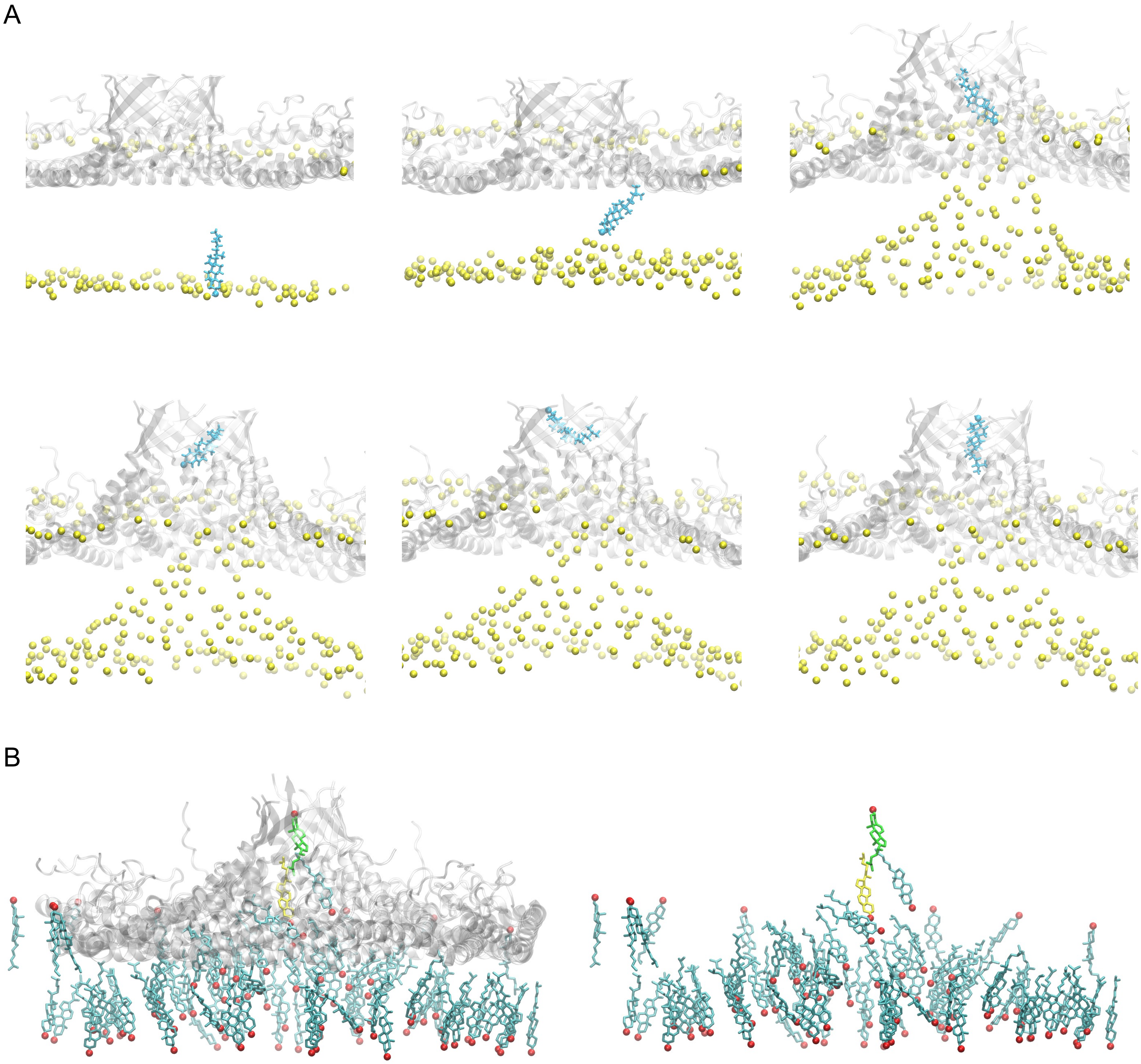


**Figure S8. Flipping of Chol into the $\beta$-barrel of CAV1-8S in all-atom simulation with a POPC/Chol membrane.** (A) Simulation snapshots illustrating the sequence of events underlying the flipping of one Chol molecule from the exoplasmic (protein-distal) leaflet into the CAV1-8S barrel in one of the replica simulations. Lipid phosphate atoms are shown as yellow spheres, protein is rendered in gray cartoon representation and the Chol molecule is shown in cyan color with its OH group as a VdW sphere and the rest of the molecule in Licorice representation. (B) Snapshots of the final frame in one of the all-atom replica simulations of CAV1-8S embedded in a POPC/Chol 70/30 bilayer. Shown are the protein in gray cartoon representation and all Chol molecules within 5 Å of protein residues in cyan Licorice (with their OH groups as red spheres). Two Chol molecules, highlighted in yellow and green, which started in the distal leaflet at the beginning of the simulation, flip into the barrel by the end of the simulations. The protein structure is hidden in the snapshot on the right to better show the conformations of Chol molecules directly interacting with the hydrophobic surface of the complex.

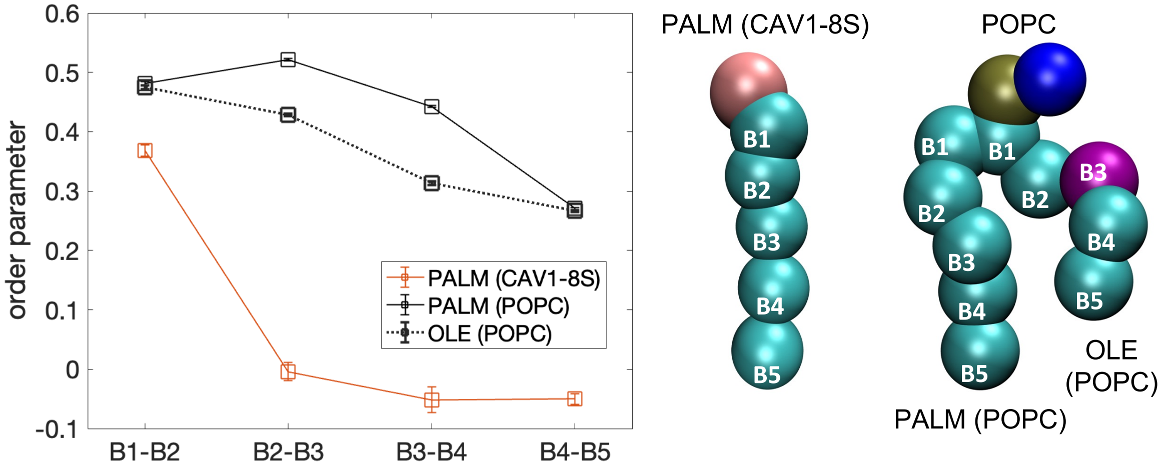


**Figure S9. Order parameters of the palmitoyl chains of CAV1-8S in the simulation with the POPC+Chol bilayer.** Shown are the P2 order parameters calculated for the bonds between consecutive beads on the palmitoyl chains as depicted on the right. Also shown for comparison are the analogous P2 order parameters of the two chains (palmitoyl, PALM, and oleoyl, OLE) of the POPC lipids in the bilayer as enumerated on the POPC schematic. The backbone bead of CAV1-8S to which the palmitoyl chain is attached is colored pink, the phosphate group of POPC is in tan and the choline headgroup is in blue.

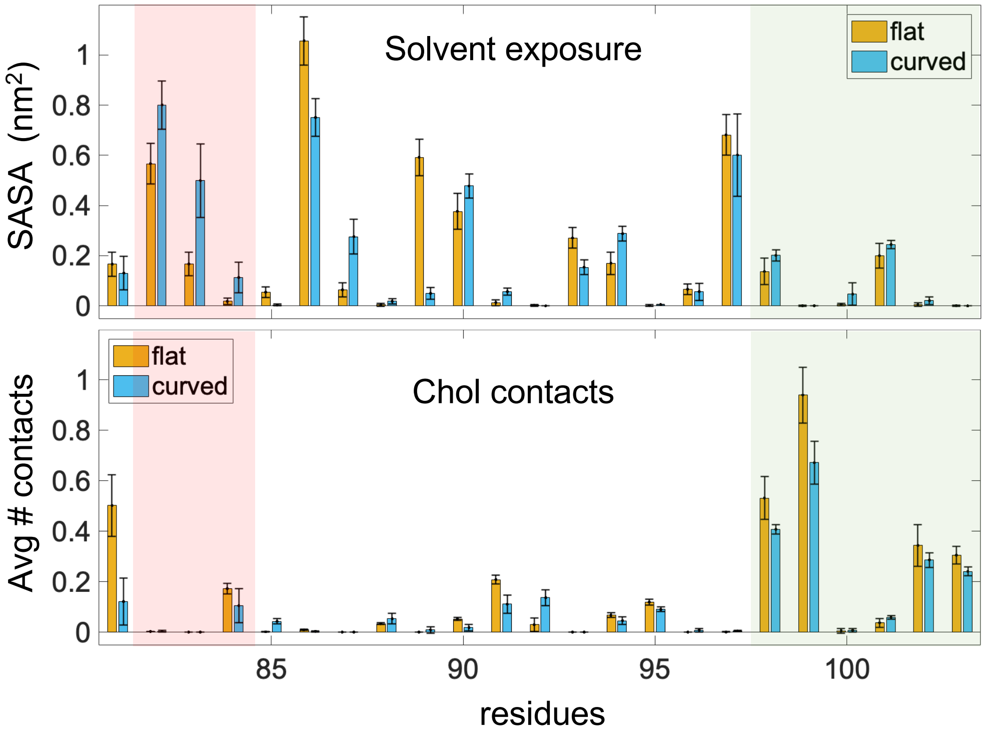


**Figure S10. Solvent exposure and Chol contacts of CAV1-8S residues in coarse-grained simulations of the symmetric inner mixture.** Aqueous solvent-accessible surface area (SASA, top) for residues 81−103 and number of contacts with Chol (bottom) averaged over the 11 caveolin protomers from the first 30 ns (flat) and last 1.5 $\mu$s (curved) of the simulations with the symmetric ***inner*** mixture. Bars represent the mean and standard deviation across three replica simulations; SD residues 82−84 shaded in red and CARC residues shaded in green, as in Figure 5.

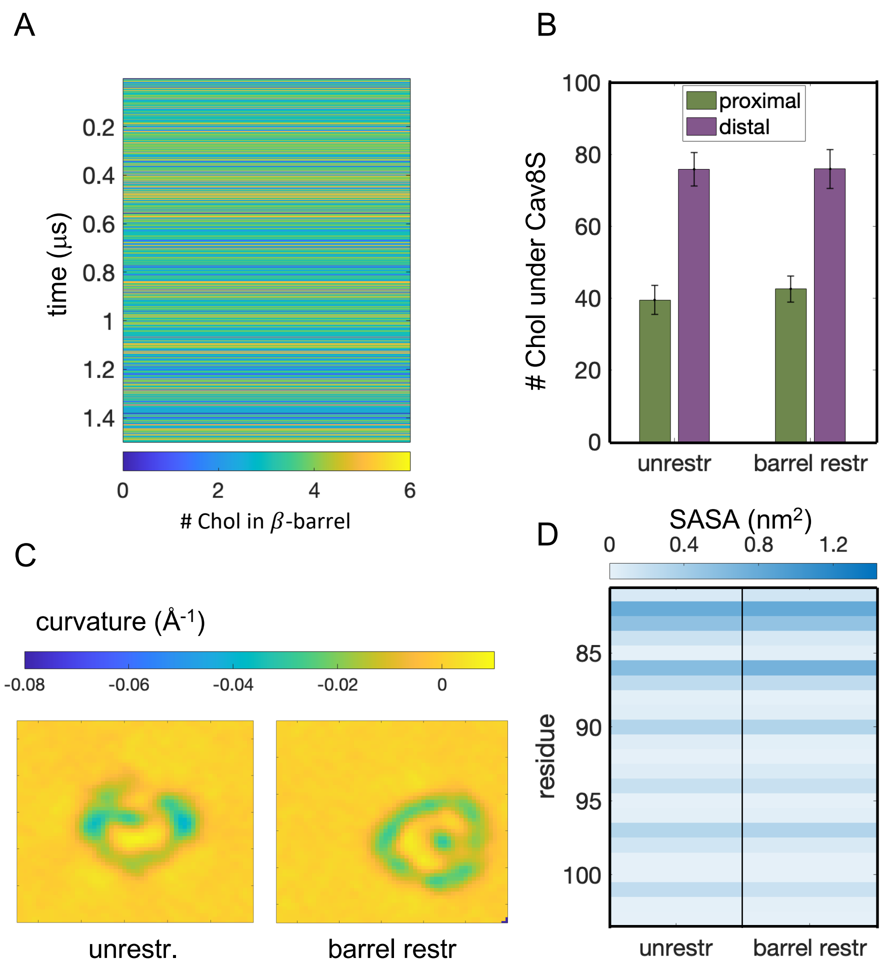


**Figure S11. Restraining the structure of the central $\beta$-barrel of CAV1-8S shows cholesterol recruitment within the barrel and does not alter the biophysical properties of the membrane.** (A) Number of Chol molecules inside the $\beta$-barrel (defined as having their headgroup ROH bead within 5 Å of the last 7 C-terminal residues of caveolin) during the last 1.5 $\mu$s of the simulation. (B) Number of Chol molecules in the proximal and distal leaflets under CAV1-8S in the equilibrium unrestrained simulation (left), and the barrel-restrained trajectory (right). Data shows the mean and standard deviation over the last 1.5 $\mu$s of the simulations. (C) Curvature maps around the complex of the two simulations, each 165 × 165 Å. Values around 0 in orange and yellow indicate regions that are flat or slightly curved away from the complex while positive values in blue represent concave morphology curved towards the protein. (D) Solvent-accessible surface areas (SASA) of residues 81−103 averaged over the 11 caveolin monomers in the 8S complex in the last 1.5 $\mu$s of the simulation trajectories.

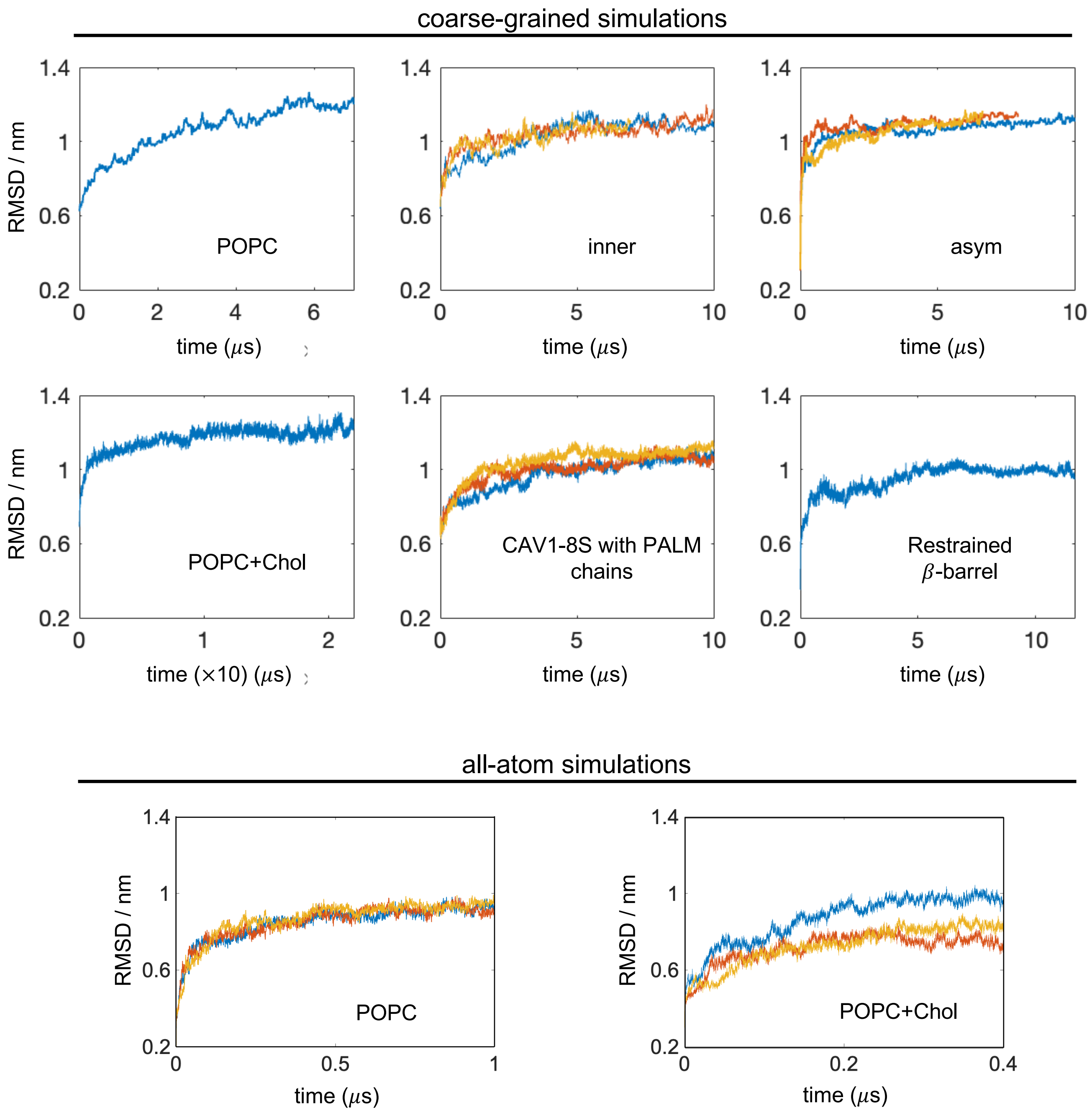


**Figure S12. Evolution of CAV1-8S structure in the single-8S simulations.** Plotted is the root mean square deviation (RMSD) from the starting structure of the complex using all backbone and side chain beads (for the coarse-grained trajectories) or all atoms (for the all-atom simulations) for the calculation. Multiple colors denote replica simulations. Note that the y-axis range is the same on all plots, but the x-axis range is different due to the different lengths of the trajectories.
